# Branch point evolution controls species-specific alternative splicing and regulates long term potentiation

**DOI:** 10.1101/2022.09.09.507289

**Authors:** Andreas Franz, A. Ioana Weber, Marco Preußner, Nicole Dimos, Alexander Stumpf, Yanlong Ji, Laura Moreno-Velasquez, Anne Voigt, Frederic Schulz, Alexander Neumann, Benno Kuropka, Ralf Kühn, Henning Urlaub, Dietmar Schmitz, Markus C. Wahl, Florian Heyd

## Abstract

Regulation and functionality of species-specific alternative splicing has remained enigmatic to the present date. Calcium/calmodulin-dependent protein kinase IIβ (CaMKIIβ) is expressed in several splice variants and plays a key role in learning and memory. Here, we identify and characterize several primate-specific *CAMK2B* splice isoforms, which show altered kinetic properties and changes in substrate specificity. Furthermore, we demonstrate that primate-specific *Camk2β* alternative splicing is achieved through branch point weakening during evolution. We show that reducing branch point and splice site strengths during evolution globally renders constitutive exons alternative, thus providing a paradigm for *cis*-directed species-specific alternative splicing regulation. Using CRISPR/Cas9 we introduced a weaker human branch point into the mouse genome, resulting in human-like *CAMK2B* splicing in the brain of mutant mice. We observe a strong impairment of long-term potentiation in CA3-CA1 synapses of mutant mice, thus connecting branch point-controlled, species-specific alternative splicing with a fundamental function in learning and memory.

## Introduction

Advances in RNA-sequencing have revealed the tremendous impact of alternative splicing on transcriptome diversity, which is especially prevalent in higher-order organism. Alternative splicing is a dynamic process that can be regulated in a tissue-, developmental-, disease-, circadian- or temperature-dependent manner (Preußner et al., 2017; Preußner et al., 2014; Ule and Blencowe, 2019). Similar to gene expression, an extensive network of *cis*-acting sequence elements and associated *trans*-acting protein factors coordinates this process and ensures its fidelity. The basic principles governing splicing regulation have been conserved across evolution, but the complexity of the spliceosome and splicing regulators has increased during the evolution of complex organisms, likely to generate the regulatory capacity for the vast amount of alternative splicing events (Ajith et al., 2016; Brooks et al., 2011; Keren et al., 2010; Ule and Blencowe, 2019; Witten and Ule, 2011). While several studies have shown that alternative splicing is controlled in a species-specific manner (Barbosa-Morais et al., 2012; Graveley, 2008; Merkin et al., 2012), the regulation and functionality of species-specific alternative splicing remains enigmatic.

Whereas gene number roughly correlates with the complexity of unicellular species, such as *Escherichia coli* or *Saccharomyces cerevisiae*, this does not hold true for higher eukaryotes. Already during early stages of the human genome project and similar efforts, it was revealed that the number of protein-coding genes in vertebrates is far below the number anticipated necessary for the phenotypic complexity. Early predictions thus suggested transcriptome diversity generated by alternative splicing to be key in creating biological complexity (Ewing and Green, 2000). In general, the frequency of alternative splicing has increased during animal evolution, with the highest frequencies detected in the primate nervous system (Barbosa-Morais *et al.*, 2012; Kim et al., 2006). This general increase in alternative splicing is strongly enriched in frame-preserving events, suggesting functional relevance (Grau-Bové et al., 2018). Additionally, alternative splicing patterns have rapidly diverged between species (Modrek and Lee, 2003; Pan et al., 2004) and are now more similar between different organs within one species, than they are between the same organs of different species (Barbosa-Morais *et al.*, 2012; Merkin *et al.*, 2012).

Species-specific splicing events appear to be largely *cis*-regulated (Barbosa-Morais *et al.*; Gao et al.), suggesting that the regulatory principles of *trans*-acting protein factors have been largely conserved during evolution. The binding codes of these splicing regulators seem largely invariant, whereas the regulatory modules and genes they affect are highly plastic and more likely to vary during evolution (Brooks *et al.*, 2011; Ule and Blencowe, 2019). These species-specific differences in splicing are not limited to animals, since similar observations have been made for various plant species (Kannan et al., 2018; Shi et al., 2019). The phenomenon is also not restricted to splicing alone: species-specific conversions of other *cis*-acting regulatory elements, such as the transition from a transcription enhancer to a promotor sequence, have also been reported (Carelli et al., 2018). Nevertheless, gene expression patterns are predominantly tissue- or organ-specific and have been largely conserved during vertebrate evolution (Barbosa-Morais *et al.*, 2012; Lin et al., 2014).

As species-specific differences in alternative splicing have been suggested to be controlled by *cis*-acting elements, the prevailing model states that this is the result of a particular combination of binding motifs of splice-regulatory proteins in the vicinity of species-specific alternative exons. However, this model falls short of explaining species-specific alternative splicing across different organs with vastly different *trans*-acting environments, leaving the mechanistic basis for species-specific alternative splicing an open question.

Few examples of species-specific alternative splicing events have been reported and analyzed in more detail. Functional consequences range from altering the activity of RNA-binding proteins (Barbosa-Morais *et al.*, 2012; Gueroussov et al., 2015) to regulating cell-cycle arrest (Sohail and Xie, 2015) or converting a noxious heat-sensitive channel into sensing infrared radiation in vampire bats (Gracheva et al., 2011). In a previous study, we have shown that the strain-specific splicing of *Camk2.1* in the marine midge *Clunio marinus* acts as a mechanism for natural adaptation of circadian timing (Kaiser et al., 2016). In vertebrates, orthologs of this gene have been identified as key regulators of neuronal plasticity and a potential species-specific regulation could thus have profound repercussions on establishing cognitive abilities in higher mammals. The calcium/calmodulin-dependent protein kinase II (CaMKII) is a unique serine/threonine protein kinase that is involved in numerous regulatory pathways (Hell, 2014). In neuronal signaling, CaMKII plays a central role in the integration of the cellular calcium influx, for example through the phosphorylation of ion channels, a key mechanism underlying synaptic plasticity (Herring and Nicoll, 2016; Hudmon and Schulman, 2002). A unique feature of the kinase is the ability to not only respond to the amplitude, but also the frequency of the activating signal. When the calcium frequency spike exceeds a characteristic threshold, the enzyme is able to adopt a calcium-independent activation state, which persists even in the absence of the activating signal (Chao et al., 2011; Meyer et al., 1992). This process is considered to be one of the fundamental mechanisms underlying long-term potentiation (LTP), which is widely seen as the molecular basis for learning and memory (Malenka and Bear, 2004).

Whereas simple organism such as *Caenorhabditis elegans* or *Drosophila melanogaster* harbor a single ancestral CaMKII gene, duplication resulted in four genes in mammals, termed *α, β, γ* and *δ* (Tombes et al., 2003). These genes and their various splicing isoforms are expressed in a tissue-specific manner, with *CAMK2A* and *CAMK2B* being the predominant isoforms in neuronal cells. Together, they are estimated to constitute up to 1% of total brain protein in rodents (Erondu and Kennedy, 1985) and are by far the most abundant proteins in postsynaptic densities (Cheng et al., 2006). Notably, conservation in CaMKII dates back to the evolutionary stage when the first synapse was thought to have formed (Ryan and Grant, 2009) and all essential features are well conserved among metazoans. Alternative splicing of the four genes leads to the expression of over 70 distinct isoforms in mammals (Sloutsky et al., 2020; Tombes *et al.*, 2003). Genetic variation has mostly been restricted to a variable linker segment that connects the N-terminal kinase domain to a C-terminal hub or association domain. Almost all mammalian splice variants are derived from alternative splicing of one of the nine alternative exons encoding this variable segment. Of the two CaMKII genes predominantly expressed in neurons, *CAMK2A* has three reported alternative splicing isoforms. On the other hand, there are eleven known *CAMK2B* isoforms generated by alternative splicing, of which up to eight have been detected in a single tissue (Sloutsky *et al.*, 2020; Tombes *et al.*, 2003). Some of these exons and their respective splice isoforms show a tissue- or developmental stage-specific regulation and have been shown to affect the subcellular localization of the enzyme, its substrate specificity, the affinity for the activator calmodulin, or other kinetic properties of the enzyme (Bayer et al., 2002; Brocke et al., 1995; GuptaRoy et al., 2000; O’Leary et al., 2006).

Here, we report the species-specific alternative splicing of three of the four CaMKII genes (*β, γ, δ*). A detailed analysis of *CAMK2B* reveals several primate-specific splice isoforms, which are generated through exclusion of exon 16. Minigene splicing assays identify an intronic regulatory sequence responsible for the primate-specific skipping of exon 16. This regulation is independent of the *trans*-acting environment, as primate-specific exon skipping is also observed in mouse cell lines. Using RNA-Seq and minigene analysis we show that weakening of the branch point (BP) sequence during evolution directs primate-specific exon 16 exclusion. Further systems-wide analyses show that weakening of core *cis*-elements required for splicing, namely the BP and the splice sites, render constitutive exons alternative during evolution. These data provide a first mechanistic understanding of how species-specific splicing patterns can be generated, also independently of the changing trans-acting environments of different tissues. Focusing on *CAMK2B,* we show that the primate-specific protein isoforms reach a higher maximal activity in *in vitro* kinase assays and display different substrate specificity. To address *in vivo* functionality of species-specific *CAMK2B* alternative splicing, we used CRISPR/Cas9 and introduced the human intronic regulatory sequence containing the weaker BP into the mouse genome, which results in a human-like *Camk2β* splicing pattern in the brain of mutant mice. Analyses of mice with humanized *Camk2β* splicing show strongly reduced long-term potentiation in CA3-CA1 hippocampal synapses. As we have not altered exonic coding regions but only intronic splicing regulatory sequences, this mouse model sets a paradigm to address the functionality of species-specific alternative splicing. Our data strongly argue for a prominent role of species-specific alternative splicing in controlling neuronal plasticity and thus species-specific cognitive abilities.

## Results

### Alternative splicing of CaMKII is species-specific

Alternative splicing of CaMKII has long been established and multiple studies have reported developmental stage- and tissue-specific splicing events (Sloutsky *et al.*, 2020; Tombes *et al.*, 2003). Differences in splicing between species are known for organisms that are evolutionary distant from humans and often feature a single ancestral *CAMK2* gene (Kaiser *et al.*, 2016; Tombes *et al.*, 2003). In vertebrate evolution, *CAMK2* genes have largely been conserved and all mammals harbor the same four genes (Figure 1A). These genes show a conserved architecture and differences mostly relate to the presence or absence of certain exons in the variable linker domain. Alternative splicing of *CAMK2* in different vertebrates has been reported, but not systematically compared (Cook et al., 2018; Rochlitz et al., 2000; Sloutsky *et al.*, 2020; Tombes *et al.*, 2003). For a detailed analysis, we performed radioactive RT-PCR with gene-specific primers on total cerebellum RNA from human and mouse (Figure S1A). Species-specific differences in the alternative splicing pattern can be seen for three of the four *CAMK2* genes (*CAMK2B, G* and *D; CAMK2A* shows no difference), revealing higher splicing complexity in humans than in mice. For further analyses we have focused on the *CAMK2B* isoform that appears to be exclusively present in human cerebellum.

We extended our analysis to include rhesus macaque (*Macaca mulatta*) and the African clawed frog (*Xenopus laevis*) (Figure 1B). For both species, the *CAMK2B* splicing pattern resembles that found in mice. All visible splice isoforms were identified by Sanger sequencing and revealed species-specific alternative splicing of *CAMK2B* exon 16 (previously also named exon IV/V (Tombes *et al.*, 2003)), whose inclusion or exclusion leads to three species-specific splice isoforms. The shortest of these, lacking exons 13 and 16 (termed Δ13,16) can easily be identified in the polyacrylamide gel. It is mostly present in humans, but as a faint band is visible for rhesus macaque as well, we refer to the exclusion of exon 16 as primate-specific. Exon 16 is furthermore the least conserved exon in the linker segment, differs in size between the CaMKII genes and, in *CAMK2G*, contains an additional splice donor site (Tombes *et al.*, 2003). It should be noted that exons 19 to 21 were not present in any of the detected *CAMK2B* isoforms, for any of the investigated species. Therefore, the full-length (FL) isoform refers to the longest isoform detected in the cerebellum. Together, these results establish species-specific alternative splicing of *Camk2β, γ* and *δ* and reveal a novel primate-specific regulation of *CAMK2B* exon 16.

**Figure 1:**
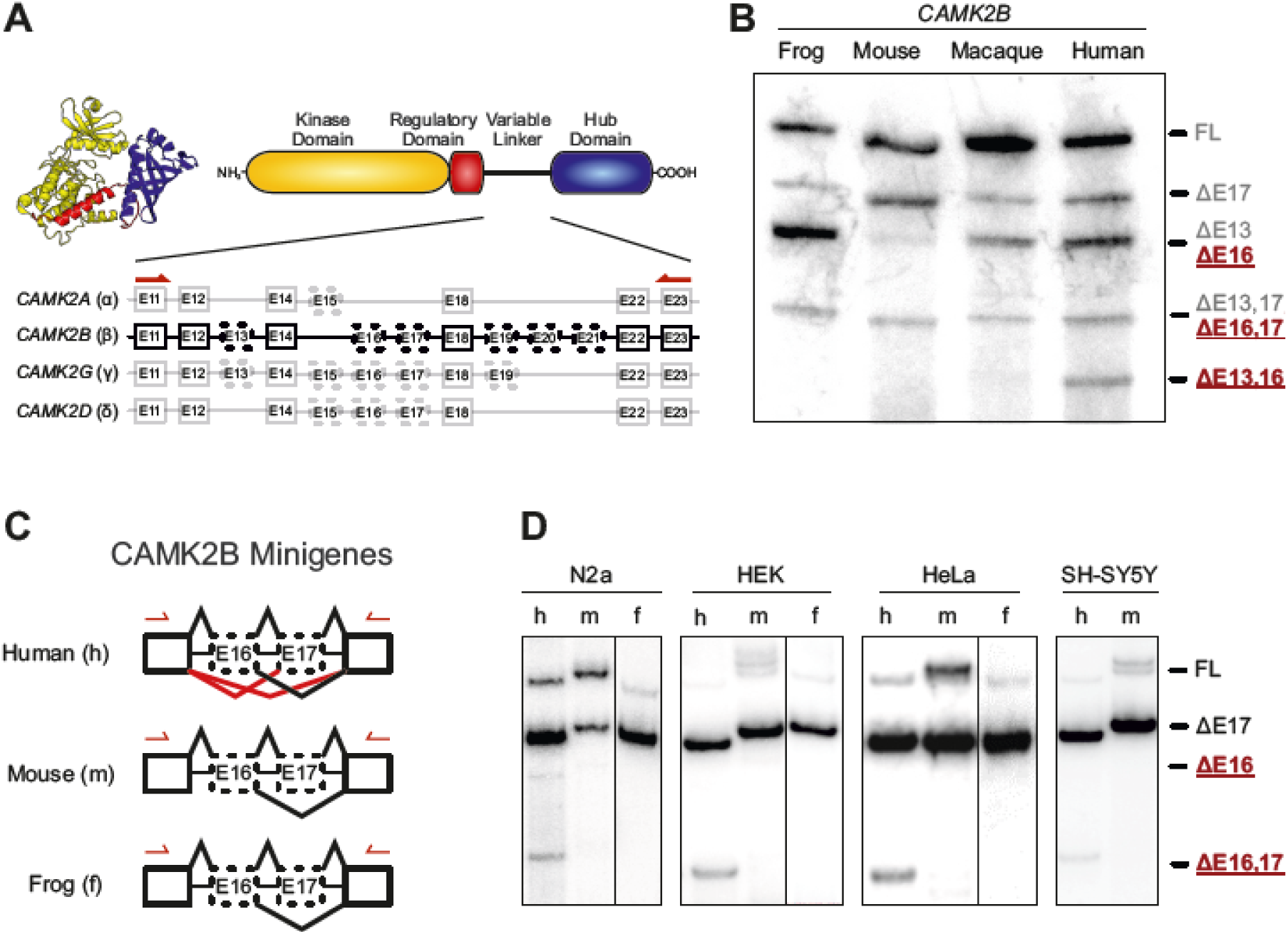
Species-specific alternative splicing of *CAMK2B* exon 16 is controlled in *cis*. (A) Schematic representation of the domain architecture of CaMKII and the intron-exon structure of the variable linker region of the four mammalian *CAMK2* genes. Numbered boxes represent exons, connecting lines represent introns. Boxes with dashed lines represent known alternatively spliced exons. (B) Endogenous *CAMK2B* splice isoforms were identified by radioactive isoform-specific RT-PCR with frog (*Xenopus laevis*), and primate (*Macaca mulatta*) total brain RNA and mouse (*Mus musculus*) and human cerebellum RNA. Isoforms were separated on a denaturing polyacrylamide gel. Isoforms are indicated on the right and named according to the exons that are skipped. As exons 19-21 are missing in neuronal tissue, they were excluded from the naming scheme. (C) Schematic representation of the minigene constructs used in D. Red lines indicate primate-specific splicing events. Arrows indicate positions of primer used for RT-PCR. (D) The human (h), mouse (m) and frog (f) (*Xenopus laevis*) sequences of exons 16 and 17, including the adjacent introns, were cloned in between two constitutive exons and transfected into N2A (mouse), HEK, HeLa and SH-SY5Y (human) cells. Resulting splice isoforms were identified by radioactive RT-PCR. Also see supplement S1.

### Species-specific *CAMK2B* alternative splicing is *cis*-regulated

Based on these findings, minigenes from human, mouse (*Mus musculus*, C57BL/6 strain) and frog (*Xenopus laevis*) were designed. The *CAMK2B* minigenes encompass two constitutive *CAMK2B* exons (exon 11 and exon 22) flanking the alternative exons 16 and 17 (Figure 1C, S1B). The introns between exons 16 and 17, and the proximal regions of the flanking introns were included as well. In order to maintain the intron/exon boundaries of the constitutive exons, the proximal region of their flanking introns was likewise inserted. The minigenes were transfected into various cell lines and the splicing patterns analyzed by radioactive RT-PCR with a vector-specific primer pair (Figure 1D). The splicing patterns of the minigenes recapitulate the observed endogenous *CAMK2B* splicing patterns. Specifically, all minigenes show bands corresponding to the full-length and Δ17 isoforms, whereas only the human minigene shows additional bands for the Δ16 and Δ16,17 isoforms. Transfection of the minigenes into various human and mouse cell lines revealed that the observed splicing pattern is independent of the cell line and species and thus of the *trans*-acting environment. This suggests a *cis*-regulated mechanism, in which differences in the pre-mRNA sequence determine the observed species-specific splicing patterns.

To pinpoint the location of the *cis*-acting element, a second set of minigenes was designed (Figure 2A). In these, intronic or exonic sequences were systematically exchanged between the human and mouse minigenes. Subsequent splicing analyses located the *cis*-acting element to the intron upstream of exon 16 (Figure 2B). Insertion of the human sequence into the mouse minigene was sufficient to induce the human splicing pattern. Conversely, transfer of the mouse sequence into the human minigene abolished exon 16 exclusion. Transfer of any other sequence did not lead to an observable change of exon 16 splicing. Together, these observations confirm the primate-specific regulation of *CAMK2B* exon 16 and show that the mechanism is *cis*-regulated, with the regulatory element located in the upstream intron.

### Branch point strength controls species-specific *CAMK2B* splicing

Having identified the approximate position of the *cis*-regulatory element, we set out to determine its exact location and sequence. As described above, the *CAMK2B* minigenes contain only a part of the intron upstream of the alternative exon 16 (Figure S1B). These 100 base pairs (bp) were further subdivided into eight overlapping segments of 20 bp (Figure 2C). The 3’ splice site itself, including the first 15 bp upstream of it, is identical between human and mouse and was thus not included in the analysis. The eight segments were exchanged between the human and mouse minigenes, and the resulting splicing patterns analyzed after expression in cell lines from both species (Figure 2D, Figure S2A). No difference between the tested human and mouse cell lines were observed, further supporting the *cis*-regulated nature of the splicing event. RT-PCR identified three segments of functional importance, two of which overlap by 10 bp. These two segments (segment 4 and 8) acted in both ways and are thus necessary and sufficient: transfer of the mouse sequence into the human minigene was sufficient to abolish the human-specific exclusion of exon 16, whereas transfer of the human sequence into the mouse context induced exclusion of exon 16. The third identified segment (segment 6) only worked in one direction: transfer from human to mouse induced exon 16 exclusion, whereas the corresponding mouse sequence inserted into the human minigene did not change the splicing pattern. Together, these findings reveal two sequences in the intron upstream of *CAMK2B* exon 16 that regulate its species-specific alternative splicing.

The presence of a *cis*-acting element suggests the existence of a corresponding *trans*-acting factor as a binding partner. In general, *cis*-acting elements act as recognition motifs for *trans*-acting proteins, which themselves are either part of the spliceosome or recruit components of it (Ule and Blencowe, 2019). We searched for candidate *trans*-acting factors by screening publicly available CLIP datasets and predicting potential binding partners based on the identified sequences (Grønning et al., 2020; Paz et al., 2014). Nine candidates, including controls, were selected and tested in an siRNA knockdown, combined with the established minigene splicing assay (Figure S2B, C), but none of the knockdowns showed a reproducible effect on exon 16 exclusion.

We therefore turned our attention to the *cis*-acting sequence itself. Notably, prediction of potential BP sequences (Corvelo et al., 2010; Nazari et al., 2019) revealed that both species harbor the most salient BP sequences in the overlap of segments 4 and 8 (Figure 2E). Strikingly, while this sequence resembles a near-optimal BP that lies within the AG dinucleotide exclusion zone (AGEZ) in the mouse intron, the corresponding human sequence scores much lower and lies slightly outside of the AGEZ. Including the other two species for which we have analyzed the endogenous splicing pattern (rhesus macaque and African clawed frog), the predicted BP strength ranks mouse > frog > rhesus > human (Table 1) and correlates well with the observed splicing pattern.

**Table 1.**
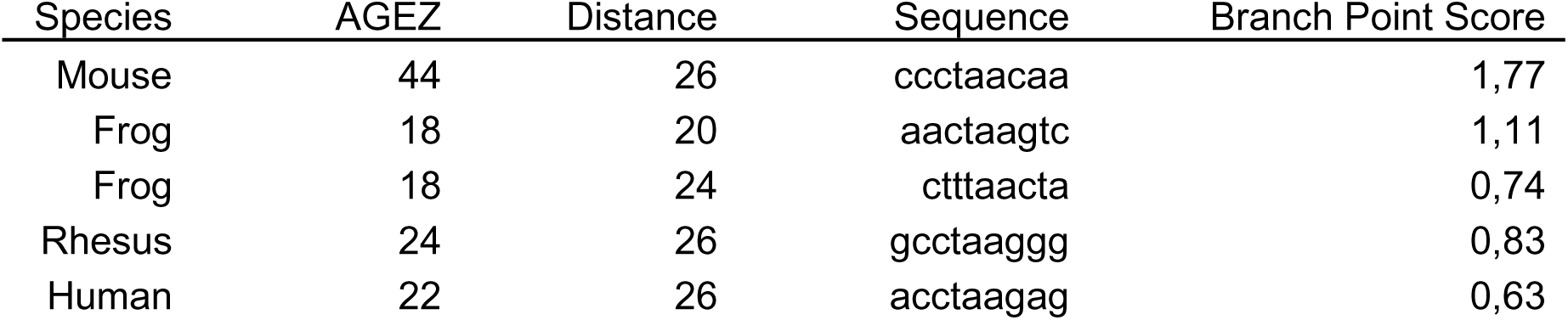
Predicted branch point scores for the intron upstream of *Camk2β* exon 16. BP scores were calculates using SVM-BP (Corvelo *et al.*, 2010). AGEZ: AG dinucleotide exclusion zone, distance: distance to 3’ splice site, sequence: sequence of identified BP, branch point score: predicted BP score (scaled vector model).

The predicted frog BP sequence scores lower than the corresponding mouse sequence, but an alternative BP is found in very close proximity. When added, the frog BP strength reaches that of the mouse sequence. The predicted rhesus sequence scores higher than the corresponding human sequence, but considerably lower than the mouse sequence. Notably, in the RT-PCR, the rhesus macaque sample also shows a faint band for the Δ13,16 exclusion isoform for endogenous *CAMK2B* (Figure 1C). We also analyzed the splice site strengths of the orthologous *CAMK2B* exons 16. As expected from the strong conservation of the splice site-proximal nucleotides, the predicted strength of the 3’ splice site does not substantially differ between mouse and human and suggests a consensus splice site in both species (MaxEntScan, human: 12.03; mouse: 13.53 (Yeo and Burge, 2004)). Similarly, the 5’ splice site is strongly conserved between both species (MaxEntScant, human: 4.41; mouse: 4.41). These data suggest that BP evolution controls species-specific alternative splicing in a way that a suboptimal BP in humans renders the exon alternative, thus creating additional *CAMK2B* complexity when compared to constitutive inclusion in mouse and frog.

To validate these finding, we designed variants of our established minigene constructs to specifically modify the predicted BP sequences (Figure 2F, G). Exchange of the 9 bp long BP motif alone was sufficient to confer species-specific splicing of exon 16 in both directions. Targeted mutation of individual nucleotides revealed that a single C to G mutation at position 7 in the mouse BP motif is sufficient to lower the predicted BP strength and induce primate-specific exon exclusion. Mutation of the BP adenine itself has a similar effect for the mouse minigene, resulting in a splicing pattern reminiscent of the human minigene. The orthogonal mutation in the human minigene has a more drastic effect, and leads to 60-80% exclusion of exon 16. This suggests the existence of additional BPs in the mouse minigene, which are absent in the human ortholog. As we had identified exchange segment 6 to be functionally relevant in the mouse sequence (Figure 2E), we exchanged a potential BP in this mouse segment to the human sequence, which does not contain a predictable BP. However, this did not alter splicing regulation, suggesting that this BP has a minor, if any, contribution to controlling exons 16 splicing in mice. Taken together, these results strongly suggest that the *cis*-regulatory element is not a conventional, splicing-factor bound enhancer or silencer motif, but that the splicing differences are instead mediated by the evolution of the BP sequence.

**Figure 2:**
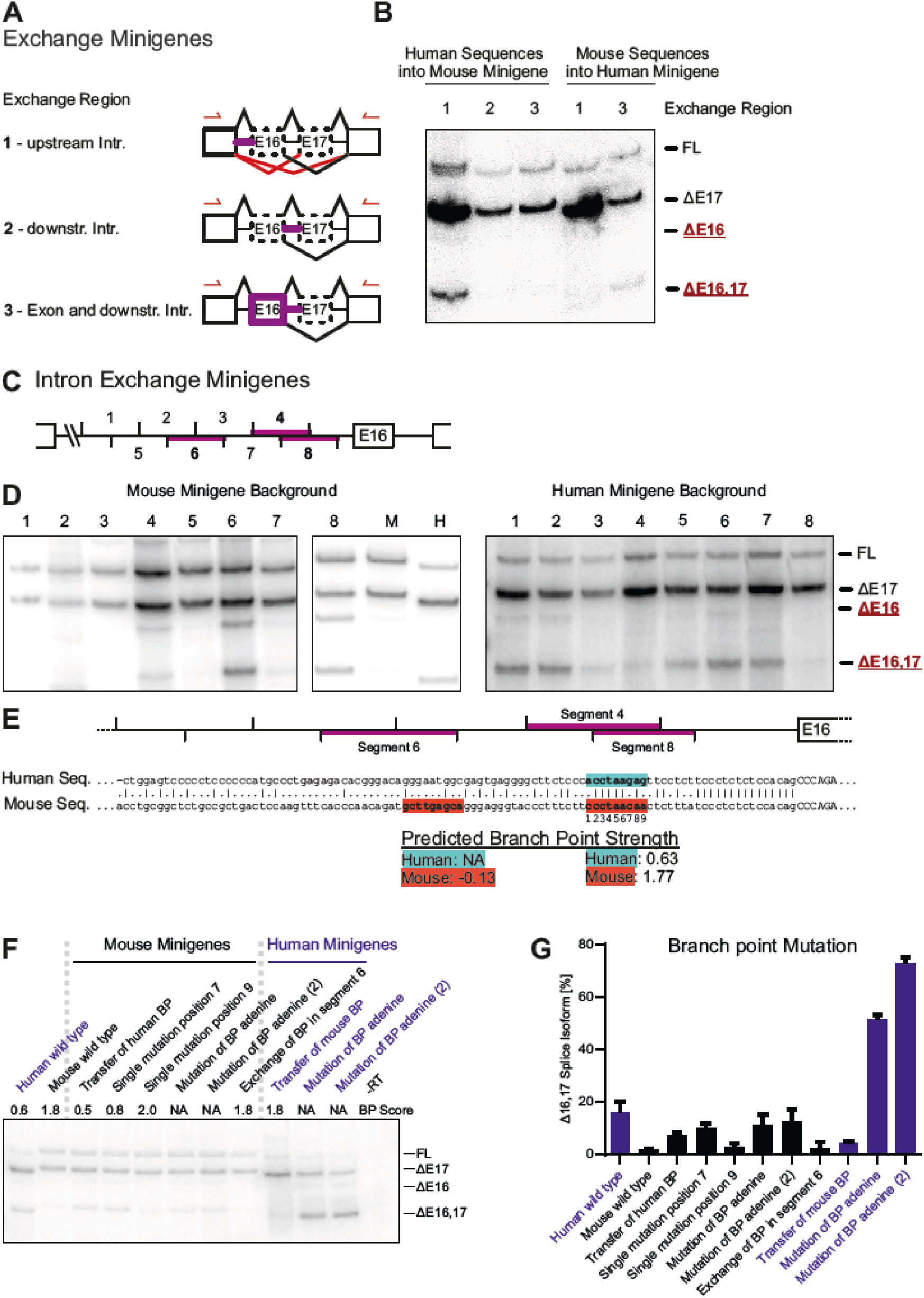
Branch point strength controls *CAMK2B* exon 16 alternative splicing. (A) Schematic representation of the minigene constructs used in B. Red lines indicate primate-specific splicing events. Purple lines highlight segments of the minigene that were exchanged between the human and mouse construct. (B) Human and mouse exchange minigenes were transfected into HEK cells and resulting splice isoforms identified by radioactive RT-PCR. n=3. (C) Schematic representation of the intron containing the identified functionally relevant *cis*-acting element. Numbers indicate 20 bp segments that were exchanged between the human and mouse construct. Purple lines highlight segments of functional relevance identified in D. (D) Human and mouse exchange minigenes were transfected into N2a cells and resulting splice isoforms identified by radioactive RT-PCR. (E) Sequence alignment between human and mouse of the intron harboring the identified *cis*-acting element. Purple lines highlight segments of functional relevance. Highlighted sequences indicate locations of predicted BPs (Corvelo *et al.*, 2010). (F) BP mutation minigenes were transfected into N2a cells and resulting splice isoforms identified by radioactive RT-PCR. (G) Quantification of F. Error bars indicate standard deviation, n=3. Also see supplement S2.

### A weak branch point correlates with *CAMK2B* exon 16 skipping across primates

To confirm these findings, publicly available RNA-Seq data from different mammals were analyzed with a focus on primates. RNA-Seq data from cerebellar tissue or, where cerebellum data were not available, total brain tissue from different species were mapped to the corresponding genome (Figure 3A). Exon 16 and exon 16,17 exclusion isoforms could be confirmed in humans, even though they only amount to ∼5-7% of all *CAMK2B* transcripts. Exclusion of exon 16 was also observed in mouse tissue, but at a much lower frequency of ∼0.4%. Even less exon 16 skipping was observed in the more distant pig (*Sus scrofa*), whereas all analyzed primates show substantial exon 16 skipping. Alignment of the BP sequences showed a clear similarity between all primates, but differences to mouse and pig (Figure 3B). The latter two species show a significantly higher BP strength, which explains the observed splicing differences. The core of the BP motif seems to be conserved among primates, with only two nucleotides showing some variation. These variations correlate with the evolutionary relationship and result in slightly different predicted BP strengths. By correlating exon 16 exclusion levels and BP strength, we observe two distinct clusters with low exon 16 exclusion and a strong BP (mouse, pig), or high exon 16 exclusion with a weak BP (primates) (Figure 3C). Even though the exact PSI (percent spliced in) values differ between human, chimpanzee (*Pan troglodytes*), bonobo (*Pan paniscus*) and gorilla (*Gorilla gorilla*), their BP sequences are identical. A recent study on the expression of CaMKII in human hippocampus found exon 16 exclusion isoforms of *CAMK2B* to range from ∼4% to 16% between tissue donors (Sloutsky *et al.*, 2020). This suggests additional regulatory layers that are specific to individual samples, for example donor age, developmental stage or the precise brain region that was used.

We then extended our analysis regarding the conservation of the BP sequence to include additional species (Figure S3A). All primates show a weak BP with a conserved sequence. This also includes the order *Dermoptera*, the flying lemurs, which are the closest relatives of primates. All other species harbor a strong BP motif, that shows a medium degree of sequence conservation among most mammals. The sequences diverge with increased evolutionary distance, but the high BP strength is maintained. Notable exceptions like, the *Anolis carolinensis* lizard, have intron sequences that do not return any valid, predicted BPs in close proximity to the splice site, suggesting fundamental differences in the splicing machinery or consensus BP sequences. Taken together, the RNA-Seq data confirm our RT-PCR analyses and minigene splicing assays, revealing species-specific differences in the BP sequence. Comparison of different mammals suggests that this feature has emerged during primate evolution and is under selective pressure, as a weak BP is maintained in all analyzed primates. The weaker BP allows alternative exon skipping to increase the diversity of *CAMK2B* transcripts and proteins, thereby controlling an essential regulator of brain development and function.

**Figure 3:**
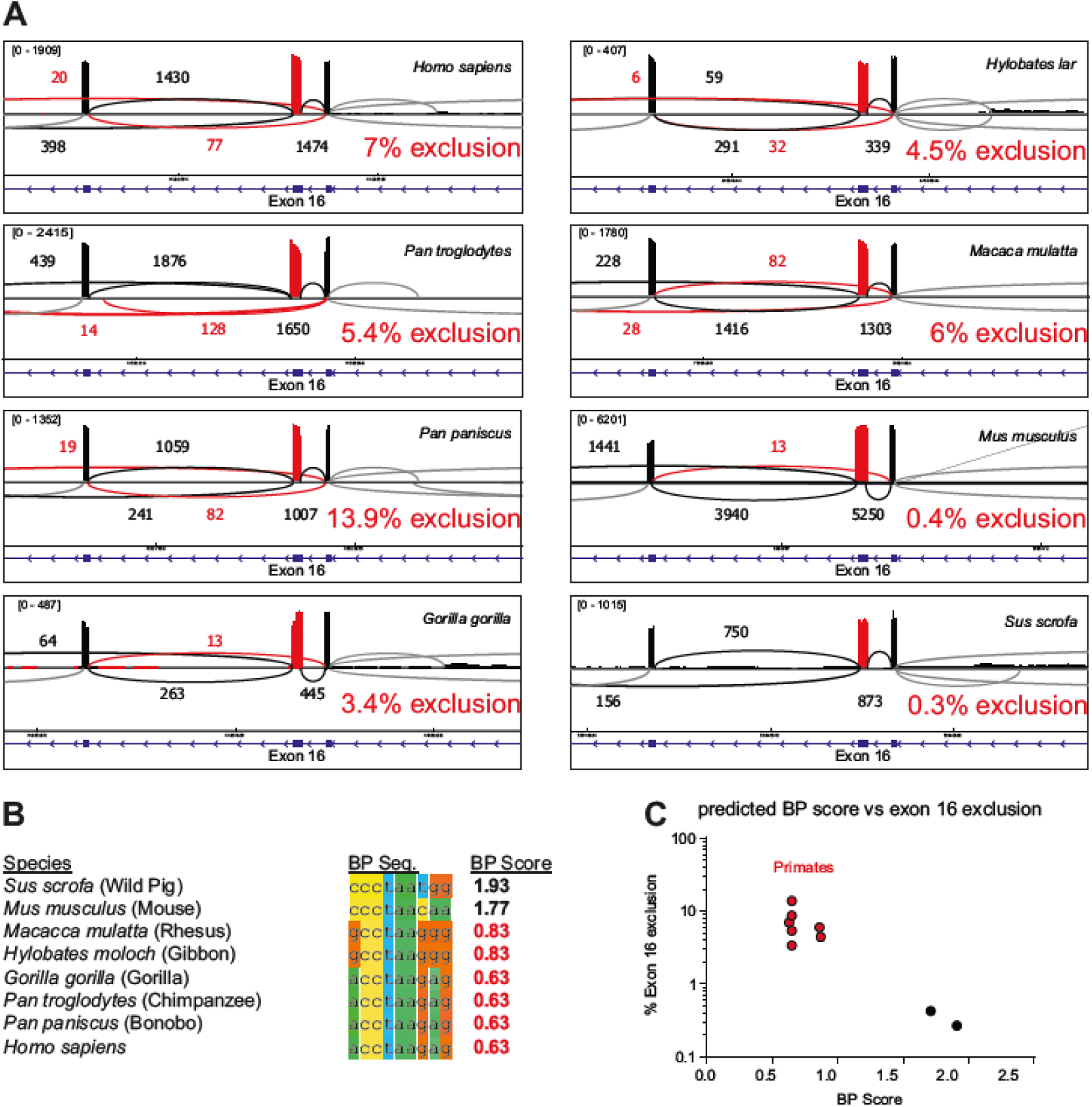
Evolutionary adaptation of branch point strength controls primate-specific *CAMK2B* exon 16 skipping. (A) Sashimi plot showing the alternative splicing of *CAMK2B* exon 16 in RNA-Seq data from human, chimpanzee (*Pan troglodytes*), bonobo (*Pan paniscus*), gorilla (*Gorilla gorilla*), orangutan (*Pongo abelii*), gibbon (*Hylobates lar*), rhesus macaque (*Macaca mulatta*), mouse (*Mus musculus*) and wild pig (*Sus scrofa*). RNA-Seq data from cerebellum was used for all species, except orangutan, for which RNA-Seq data from total brain tissue was used. Red color indicates exon 16 and exon 16 exclusion reads. Numbers indicate number of reads per splice junction, with the minimum set to 3 junction reads. Shown in blue is the intron/ exon-structure of the displayed region. % exon 16 exclusion is indicated. Splicing was analyzed from RNA Seq data using rMATS (Shen et al., 2014). (B) Alignment of the identified functionally relevant BP sequence. The BP strength was predicted using SVM-BPfinder (Corvelo *et al.*, 2010) with the human BP model. BP score refers to the BP motif score (scaled vector model). (C) The predicted BP strength from B was plotted against the exon 16 exclusion levels determined by RNA-Seq. Also see supplement S3.

### Branch point strength globally controls species-specific alternative splicing

We next addressed whether species-specific differences in the BP motifs globally regulate species-specific alternative splicing. To this end, we first defined orthologous exons between mouse and human (see Material and Methods) and then analyzed RNA-Seq data from a large collection of human and mouse brain samples. This approach allowed us to define orthologous exons that are alternatively spliced in both species, or that are alternative exclusively in mouse or human brain (Supplementary data 1). In line with the notion of an increased frequency of alternative splicing in more complex organisms, we observed a higher number of exons that are alternative only in humans (Figure 4A, B S4A, B). We then analyzed, these species-exclusive subsets of alternative exons and observed clear differences in the strengths of their core splicing elements (Figure 4C, D). Exons that are exclusively alternative in humans show a weaker BP sequence score and BP motif score (which includes the distance to the 3’ splice site), as well as weaker 3’ and 5’ splice site scores (Figure 4C) when compared to the constitutive mouse orthologs. A similar trend can be observed for mouse-exclusive alternative exons, which show reduced BP and splice site scores when compared to the constitutive human orthologs (Figure 4D). Importantly, this effect is restricted to the alternatively spliced exon itself, and the core splicing elements of the surrounding constitutive exons do predominantly not show significant differences between both species (Figure S4C), demonstrating specificity for the alternative exons. These data strongly suggest a mechanistic basis for establishing global species-specific splicing patterns through evolution of the core splicing sequences. Weakening of splice site and/or BP sequences allows alternative usage of an exon, thus increasing transcriptome and proteome complexity through suboptimal exon recognition. Notably, this model also provides an explanation for species-specific alternative splicing that is at least partially independent of the different *trans*-acting environments in different cell types and organs.

The impact of these core splicing sequences on alternative splicing is further underlined by a clear and significant correlation between the rate of exon skipping (PSI) and the strength of the splice sites and the BP when considering all alternative exons in human or mouse brain (Figure 4E). While combining BP and splice site scores in a single value (see Material and Methods) further increases the correlation with the PSI of an exon (Figure 4E), the correlation coefficients show that additional variables influence splicing outcome. These observations strongly support the conclusion that species-specific alternative splicing is, to a large part, controlled through the evolution of core splicing sequences, namely the splice site and BP sequences.

**Figure 4:**
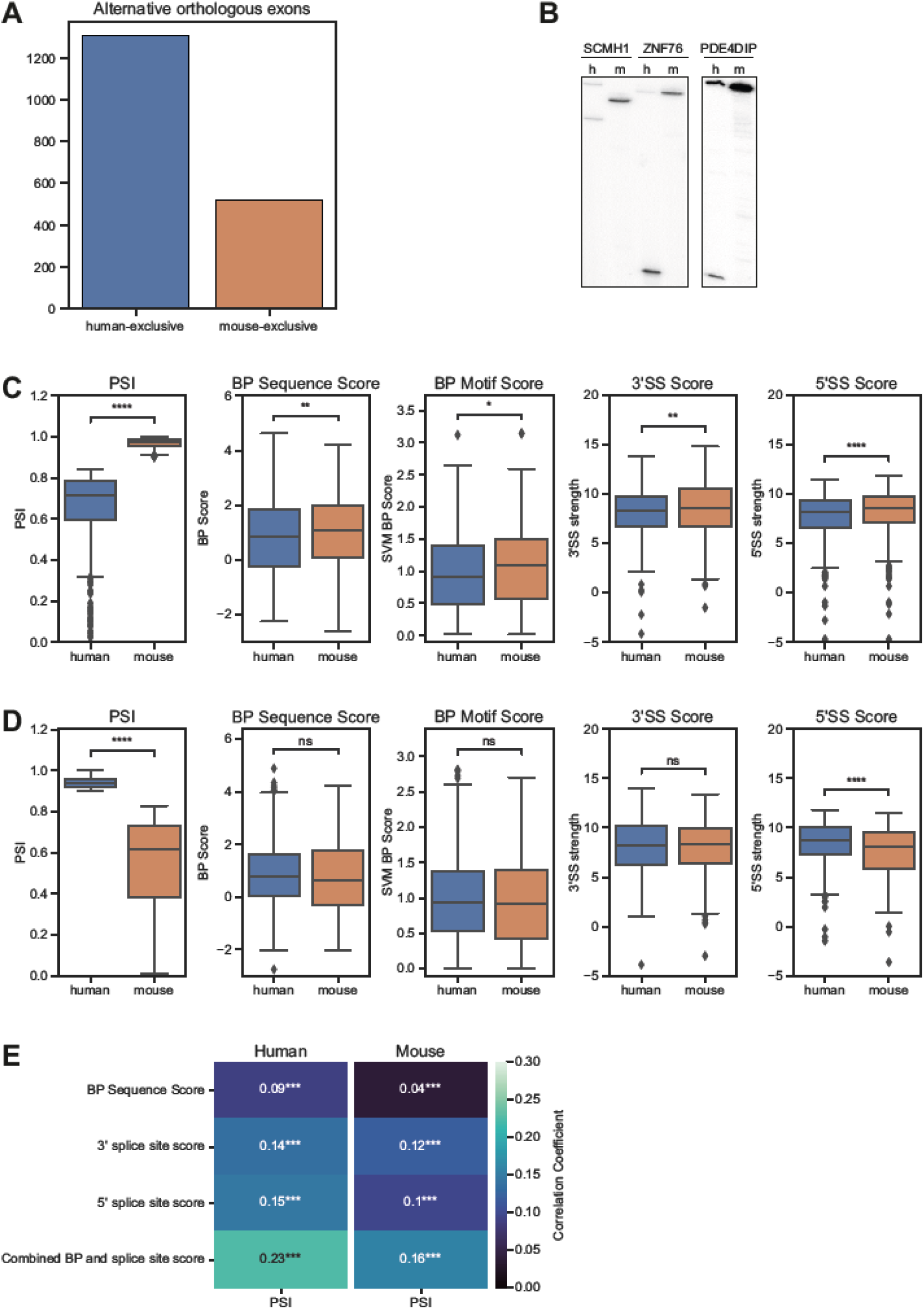
Branch point and splice site strength globally control species-specific alternative splicing. (A) Species-exclusive alternative orthologous exons. RNA-Seq data from different brain regions from mouse (n=47) and human (n=9) was analyzed to identify species-specific splicing pattern. The analysis was restricted to orthologous exons (see Material and Methods for details) that are alternatively spliced in one species (PSI < 0.9) but not the other (PSI > 0.9) (see Supplementary data 1). (B) Validation of species-exclusive alternative exons by radioactive RT-PCR. m: mouse, h: human. (C, D) Boxplots comparing human and mouse splicing element scores for human-exclusive (C) or mouse-exclusive (D) alternative orthologous exons. PSI: percent spliced in, BP Sequence Score: branch point sequence score, BP Motif Score: branch point motif score using a scaled vector model (Corvelo *et al.*, 2010), 3’/5’SS Score: splice site score (Yeo and Burge, 2004). *p<0.05, **p<0.01, ***p<0.001 (paired Wilcoxon test). (E) Heatmap displaying the Spearman correlation coefficients between the PSI (percent spliced in) of all alternative exons in mouse or human brain and other parameters. RNA-Seq data from human (n=4) and mouse (n=4) cerebellum was analyzed and not restricted to orthogonal exons. Also see supplement S4. Asterisks indicate significance levels: ***p<0.001.

### Primate-specific CaMK2β isoforms display slightly increased activity

Having established the genomic causes and the transcriptomic consequences of the species-specific alternative splicing of *CAMK2B*, we set out to determine its effect on the protein level. We selected two species-specific isoforms (Δ16,17 and Δ13,16) as well as two control isoforms (FL and Δ13) for recombinant production and purification from insect cells (Figure 5A). Again, the full-length isoform refers to the longest detected isoform in cerebellum and lacks exons 19 to 21 (Figure 1A, B). These four isoforms were tested in a radioactive *in vitro* kinase assay with the model substrate Syntide 2 (Hashimoto and Soderling, 1987), linked to GST (Figure 5B, C). Activity was monitored as a function of calmodulin concentration to test the cooperativity of the enzyme. Consistent with a recent publication (Sloutsky *et al.*, 2020), we did not observe major differences in the EC_50_ values or the Hill coefficients (Table 2) between the four CaMKIIβ variants. Instead, we observed small but significant differences in the maximal activity (V_max_) reached at optimal calmodulin concentrations (Figure 5D). At concentrations of 100-1000 nM calmodulin, both primate-specific protein isoforms reach a slightly higher maximal activity compared the FL and Δ13 isoforms. The same effect was also observed using human full-length tau protein (tau 441) as an alternative CaMKIIβ substrate (Figure S5A, B).

One of the key properties of CaMKII is its ability to adopt different activation states, based on its own phosphorylation pattern (Bayer and Schulman, 2019). Upon stimulation, the enzyme quickly *trans*-autophosphorylates on T287 and adopts an auto-activated state, that persists even in the absence of calcium/calmodulin. Recent studies suggest that the rate at which certain activating and inhibiting phosphorylations are acquired differs between CaMKII protein isoforms and might also be influenced by the length and composition of the variable linker segment (Bhattacharyya et al., 2020). We thus tested the autoactivity of the selected CaMKIIβ isoforms at varying calmodulin concentrations, which directly reflects the activation and hence phosphorylation state of the enzyme. CaMKII was first stimulated in the absence of the substrate protein, after which calcium was quenched by adding EGTA. Addition of the substrate Syntide 2-GST allowed assessment of the pre-established autoactivity previously generated. Similar differences regarding V_max_ could be observed, with the primate-specific protein isoforms reaching slightly higher maximal activities (Figure S5C, D). Together, these results confirm that the tested CaMKIIβ splice isoforms do not differ in their EC_50_ values or Hill coefficients. Instead, a slight difference in maximal activity at optimal calcium/calmodulin concentrations can be observed. Notably, this may allow the primate-specific variants to react more strongly to calcium influx and may thus contribute to translate primate-specific alternative splicing into functionality.

### CaMKIIβ isoforms have different substrate spectra

In addition to subtle kinetic variations between the CaMKIIβ isoforms, we considered differences in their substrate spectra as a further mechanism for diversified functionality. Instead of testing individual substrates *in vitro*, which has previously been done for fly CaMKII (GuptaRoy *et al.*, 2000), we adopted the analog-sensitive kinase system (Lopez et al., 2014). This approach allows for direct labelling of kinase substrates in complex samples and does not require prior knowledge of potential phosphorylation targets.

An analog-sensitive variant has previously been described for CaMKIIα (Wang et al., 2003) and, consistent with high sequence similarity of the kinase domains, the same residue exchange (F89G) was effective in creating a CaMKIIβ variant that could use ATP analogs with bulky side chains on their N^6^-atoms (Lopez *et al.*, 2014). We confirmed *in vitro* and in cells that the analog-sensitive variant exhibited similar enzymatic activity as the wt enzyme, that only the variant could be competitively inhibited by bulky ATP analogs and that in permeabilized N2A cells, the ATP analog N^6^-benzyl-ATPγS was used only by the variant kinase (Figure S6A-E).

The four CaMKIIβ isoforms that were analyzed in *in vitro* kinase assays and two additional control samples - untransfected (UT) and a kinase dead variant (K43R) - were chosen for kinase assays with N^6^-benzyl-ATPγS in permeabilized N2A cells, subsequent thiophospho-enrichment and detection of substrates *via* mass spectrometry (MS) analysis (Supplementary data 2). To identify qualitative and quantitative differences in the substrate spectra, we excluded targets also identified in the control samples, or mapping to the alternative exons in the variable linker itself. We then generated a correlation matrix based on the abundance of the identified phosphorylation sites. Strong correlations between the FL and Δ16,17 isoforms on the one hand and between the Δ13 and Δ13,16 isoforms on the other were observed, indicating that different splice isoforms have preferred substrates (Figure 5E). Interestingly, the correlation between the FL and Δ16,17 isoforms is mainly based on similar CaMKII autophosphorylation (Figure S6G, Table S1), which likely controls kinase activity and/or localization. Comparing substrates that are phosphorylated by the different variants also identified substrates that are exclusively phosphorylated by individual isoforms, including 17 targets of the primate specific Δ13,16 isoform. We also note that the largest intersection is between all four CaMKIIβ isoforms, indicating a relatively large overlap in their substrate spectra (Figure 5F). While we did not observe clear-cut differences in the gene ontology (GO) terms of isoform-specific substrates, our data strongly suggests that CaMKIIβ isoforms have isoform-preferred/specific substrates. For example, substrates only found for the Δ16,17 isoform are enriched in the GO term “myelin sheath” (GO:0043209) (Figure 5G), suggesting a potential isoform-specific functionality. Interestingly, a substrate specifically phosphorylated by the primate-specific isoforms is the catalytically-relevant Y-box of phospholipase C β1 (Plcb1) (Supplementary data 2), an enzyme involved in inositol triphosphate (IP_3_) signaling that has been connected to learning and memory (Cabana-Domínguez et al., 2021). These results suggest that apart from differences in catalytic activity, CaMKIIβ isoforms differ in their substrate preferences, with primate-specific isoforms preferentially targeting specific proteins related to neuronal functions.

**Figure 5:**
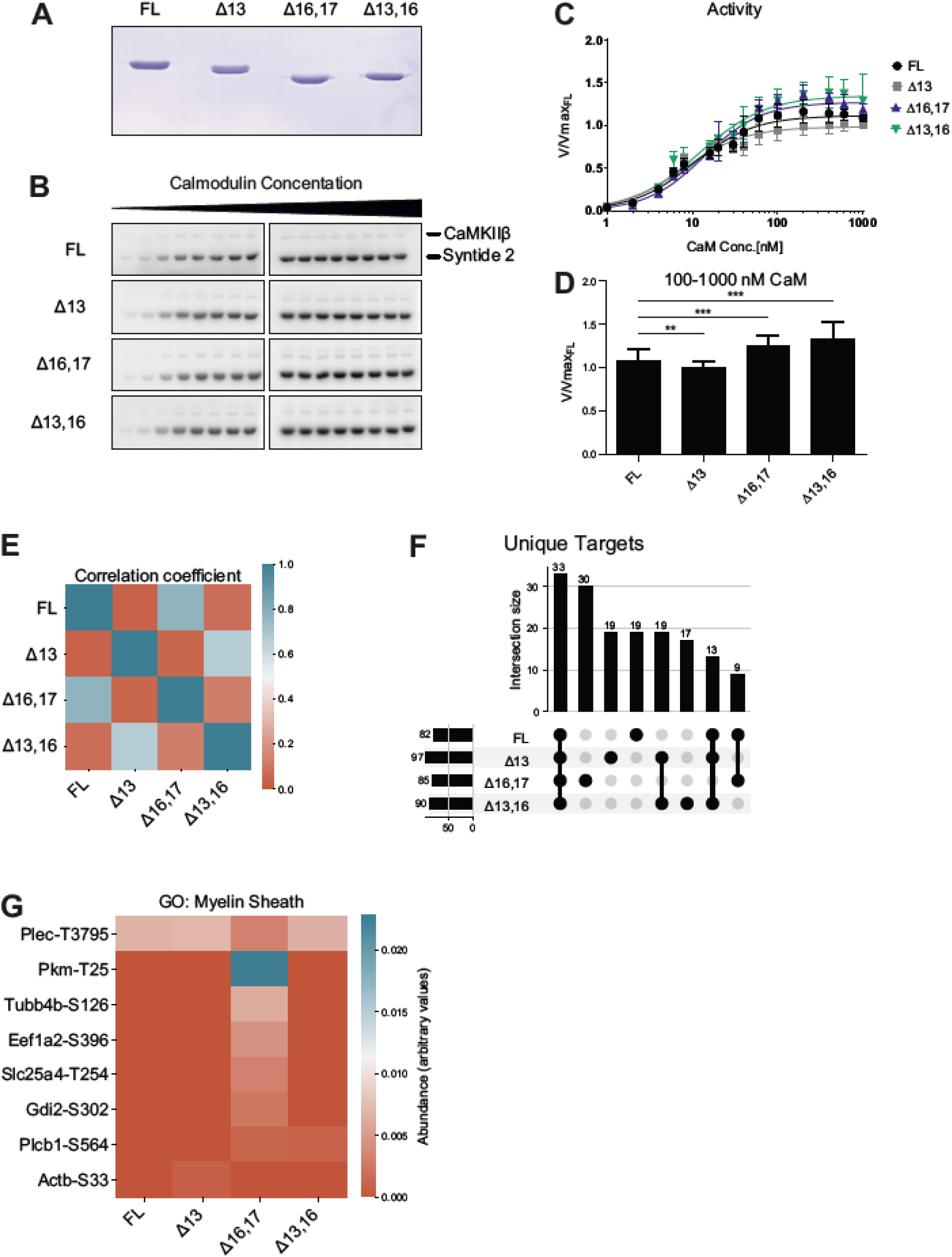
CaMKIIβ protein isoforms differ in their kinetic properties and substrate spectra. (A) SDS-PAGE of purified CaMKIIβ isoforms. Proteins were expressed in insect cells and purified via Strep-affinity and size exclusion chromatography. Protein concentration was determined via UV-absorption at 280 nm and precisely levelled by repeated SDS-PAGE, Coomassie-staining and subsequent quantification. (B) *In vitro* kinase assay with different CaMKIIβ isoforms. CaMKII activity against a protein substrate (Syntide 2, fused to GST) was measured as a function of calmodulin concentration. Direct phosphorylation of the substrate by CaMKIIβ was measured via ^32^P incorporation. Samples were separated on an SDS-PAGE and detected using autoradiography. (C, D) Quantification of B, normalized to the maximum activity of the FL isoform (n = 6). Error bars indicate standard deviation. Data was fitted to a Hill equation. (D) Samples at maximal activity were combined. Error bars indicate standard deviation. *p<0.05, **p<0.01, ***p<0.001 calculated by Student‘s or Welch‘s t-test and adjusted for multiple comparisons using Holm‘s method. (E) Correlation matrix of the substrate spectra of different CaMKIIβ isoforms, as determined by an analog-sensitive kinase assay. The analysis was restricted to CaMKIIβ-specific targets. A Person correlation coefficient was calculated based on the intensity values of individual phosphorylation sites. (F) Intersection plot showing the isoform and group-exclusive phosphorylation sites. Analysis was restricted to CaMKIIβ-specific targets. Numbers on the left indicate the total number of phosphorylation sites detected in a sample. Numbers on the top indicate the intersection size between samples, meaning the number of phosphorylation sites that are unique to this group of samples. Black dots and connecting lines indicate the exact group of samples for which the intersection size is displayed (G) Heatmap showing the abundance of individual phosphorylation sites associated with the GO term “myelin sheath” (GO:0043209) in the substrate spectra of different CaMKIIβ isoforms. Also see supplement S5 and S6.

**Table 2.** Kinetic parameters of purified CaMKIIβ isoforms. Kinetic parameters as determined by in vitro kinase assay and subsequent fitting of a Hill equation. h: Hill Coefficient.

|  | Substrate: Syntide 2 |  |  |  | Substrate: Tau-441 |  |
| --- | --- | --- | --- | --- | --- | --- |
|  | Isoform |  |  |  | Isoform |  |
| | FL | $\Delta 13$ | $\Delta 16,17$ | $\Delta 13,16$ | FL | $\Delta 16,17$ |
| $V_{\max}$ | 1.11 ± 0.02 | 0.99 ± 0.02 | 1.27 ± 0.03 | 1.35 ± 0.03 | 1.04 ± 0.02 | 1.25 ± 0.03 |
| $h$ | 1.22 ± 0.11 | 1.20 ± 0.09 | 1.29 ± 0.11 | 1.11 ± 0.11 | 1.53 ± 0.19 | 1.58 ± 0.17 |
| $EC_{50}$ | 10.30 ± 0.82 | 7.80 ± 0.56 | 13.65 ± 0.97 | 11.98 ± 1.16 | 12.2 ± 0.90 | 18.8 ± 1.30 |

### A mouse model with humanized *Camk2β* splicing

To study the consequences of *CAMK2B* alternative splicing *in vivo*, we generated a mouse model with humanized *Camk2β* splicing pattern. Based on the results obtained with our minigenes, we used CRISPR/Cas9 and introduced the identified human intronic regulatory sequence, including the BP, into the mouse genome. We generated two mutant mouse strains, one containing the human intronic regulatory sequence, termed “humanized strain”, and one in which the mouse sequence had simply been deleted, termed “deletion strain” (Figure 6A). The intron-exon structure was retained for both strains, as only a part of the intron was exchanged or deleted, leaving all splice sites intact. Both strategies led to a human-like *Camk2β* splicing pattern in the brain of the mutant mice, revealed in particular by the emergence of the primate-specific splice isoform Δ13,16 (Figure 6B). Sanger sequencing also confirmed the presence of the other previously identified primate-specific exon 16-exclusion isoforms. Interestingly, both mouse strains showed an additional band for a Δ13,16,17 isoform, that we previously did not detect in human cells. These findings confirm the results from the minigene splicing assays and the postulated model of species-specific differences in BP strength. Furthermore, they corroborate that primate-specific *Camk2β* splicing is *cis*-regulated, with the mouse sequence harboring a functionally relevant, strong BP motif. The sequence of the human intron contains a weak BP, and its knock-in into the mouse genome has a similar effect as simply deleting the strong mouse BP.

There are clear differences in the splicing patterns of heterozygous and homozygous animals. Whereas the heterozygous animals showed a human-like *Camk2β* splicing pattern, the homozygous animals lacked the FL and Δ17 isoforms. Instead, exon 16 was efficiently skipped in these animals, as revealed by the strong presence of the Δ16 and Δ16,17 isoforms. To confirm these observations, we performed RNA-Seq on cerebellum samples from the humanized strain. This analysis showed almost 100% inclusion of exon 16 in the wild type mouse, which was reduced to around 50% in heterozygous animals and essentially absent in homozygous animals (Figure 6C). We also checked whether alternative splicing was affected on a global level in the humanized mouse strain of our mouse model. However, only exon 16 of *Camk2β* was found to be substantially and significantly differentially spliced (Figure 6D), suggesting that alternative splicing is not globally affected in the humanized mouse strain. We also did not detect any significant differences in gene expression levels between wild type and heterozygous animals, and only minor differences between wild type and homozygous animals (Figure 6E). These results suggest that under resting conditions, neither global gene expression nor global alternative splicing are significantly altered in our humanized *Camk2β* mouse model.

### Mice with humanized *Camk2β* splicing pattern show reduced levels of LTP

Having confirmed the validity of our *in vivo* model, we next set out to determine the effects of altered *Camk2β* splicing on synaptic plasticity. We performed an electrophysiological characterization of CA3-CA1 synapses in acute hippocampal slices of homozygous humanized or BP-deleted strains. Basal synaptic transmission, as well as short-term plasticity, measured as paired pulse ratio (PPR), were unaltered in the mutant mice (Figure S7A, B, mean ± SD wt: 1.33 ± 0.15, humanized 1.29 ± 0.21; deletion 1.27 ± 0.09). However, high frequency-induced long-term potentiation (LTP) was significantly impaired in both mouse strains, 30 minutes post stimulation (Figure 6F-H, normalized amplitude wt: 1.285 ± 0.20, humanized 1.09 ± 0.14; deletion 1.07 ± 0.12). Induction of LTP with a single high frequency tetanic pulse or with multiple pulses led to similar results (Figure S7 C-E). In contrast, short-term potentiation measured as the immediate response after the tetanic pulse (post-tetanic potentiation) was not affected (mean ± SD: wt: 2.11 ± 0.53, humanized 2.10 ± 0.29; deletion 2.18 ± 0.41). Together, these observations show that in our mouse model with humanized *Camk2β* alternative splicing basal synaptic transmission as well as short-term plasticity are unaffected, whereas LTP is severely impaired. Thus, *Camk2β* species-specific alternative splicing correlates with differential species-specific control of LTP. As we did not alter coding sequences but only replaced an intronic splicing-regulatory element, our data provide clear evidence for a prominent role of species-specific alternative splicing in controlling synaptic plasticity, which forms the molecular basis for cognitive functions.

**Figure 6:**
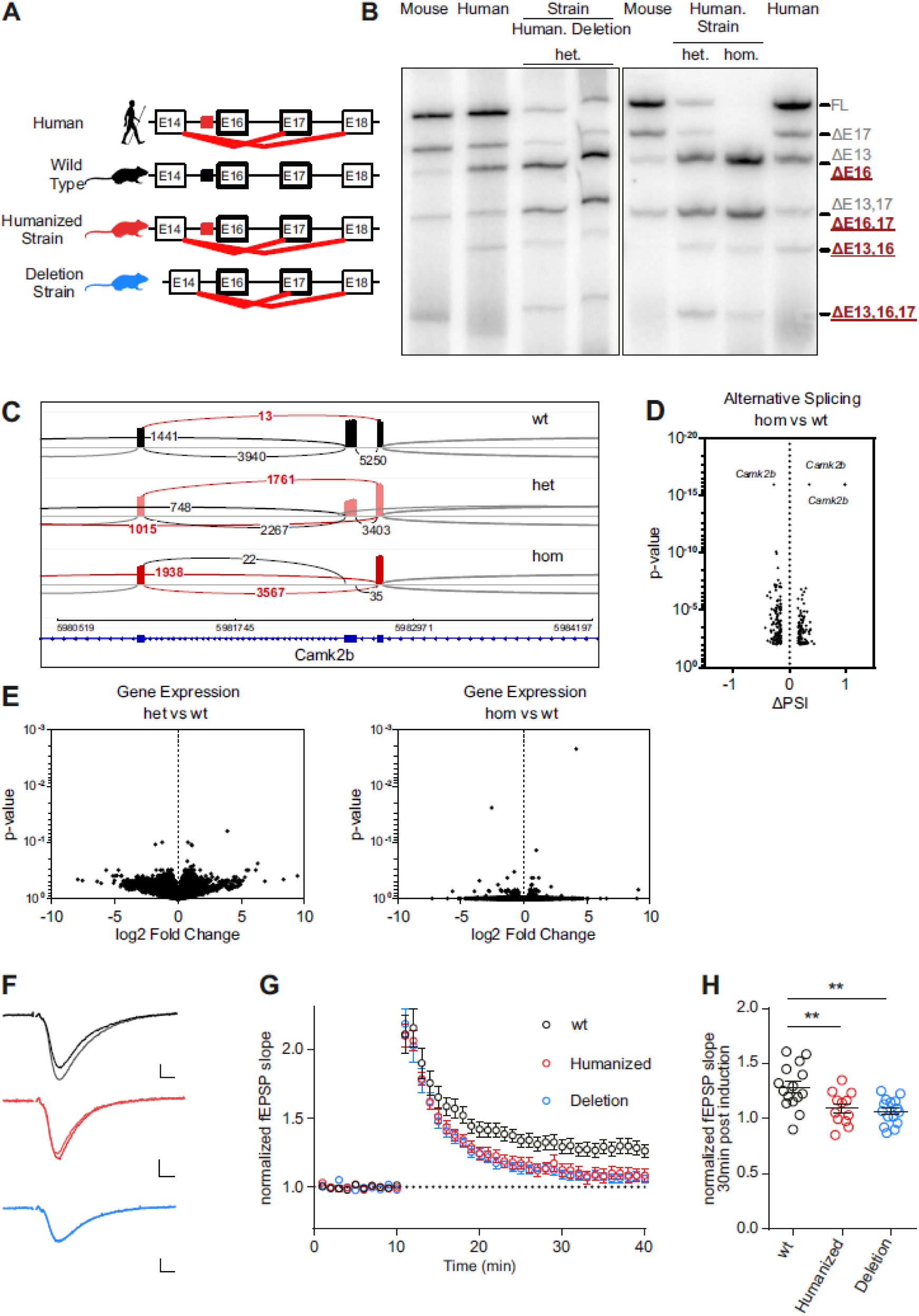
A mouse model with a humanized *Camk2β* splicing pattern shows strong impairment in LTP formation. (A) Schematic representation of the intron-exon structure of the variable linker region of the *CAMK2B* gene and comparison of the identified alternative splicing isoforms in human, wild type mice, and the novel mouse model with a humanized *Camk2β* splicing pattern (humanized strain and deletion strain). Red lines indicate identified species-specific splicing events. Colored boxes indicate the location of the identified *cis*-regulatory element in human (red) and mice (black). (B) Endogenous *Camk2β* splice isoforms were identified by radioactive isoform-specific RT-PCR with mouse (*Mus musculus*) and human cerebellum RNA. Isoforms were separated on a denaturing polyacrylamide gel. Isoforms are indicated on the right and named according to the skipped exons. Human.: humanized strain, Deletion: deletion strain, wt: wild type animals, het: heterozygous animals, hom: homozygous animals. (C) Sashimi plot showing the alternative splicing of *Camk2β* exon 16 in RNA-Seq data from wild type, heterozygous and homozygous mice of the humanized strain. Each graph summarizes RNA-Seq data of 4 biological replicates. (D) Volcano plot mapping the differences in percentage spliced in (PSI) of cassette exons of homozygous vs. wt animals against their respective p-values. Individual splicing events affecting *Camk2β* exon 16 are labelled. (E) Volcano plot mapping gene expression changes in the mouse model for both heterozygous and homozygous animals of the humanized strain against their respective p-values. (F) Example traces showing average of baseline and potentiated field excitatory postsynaptic potentials (fEPSP) 30 min after LTP induction. Scale bar: 0.2 mV/ 5 ms. (G) Time course of LTP induction in CA3-CA1 synapses in acute hippocampal slices. LTP was induced after 10 min with a single train of 100 Hz, 1 s wt (wild type): 15 slices, 6 mice, humanized (humanized strain, homozygote): 12 slices 6 mice; deletion (deletion strain, homozygote):15 slices, 6 mice. (H) Dot-plots depicting the field EPSP slope 30 min after LTP induction. **p<0.01, calculated by ANOVA followed by Dunnet’s test. Also see supplement S7.

## Discussion

Species-specific alternative splicing has been suggested to contribute to shaping species-specific properties and abilities, including cognition. However, how species-specific alternative splicing patterns are established remains enigmatic. How they translate into species-specific functionality at the level of protein isoforms, cells and the whole organism, is another fundamental, largely unanswered question that affect our very identity as humans. Here, we uncover a pervasive mechanism underlying species-specific alternative splicing, *i.e.* the species-specific degree of deviation of splice sites and, in particular, BP sequences from consensus motifs. Furthermore, to our knowledge this is the first report of species-specific alternative splicing of any mammalian CaMKII gene, *i.e.* genes that give rise to one of the most important groups of proteins shaping neuronal functions. We also demonstrate that species-specific *CAMK2B* alternative splicing is controlled by the principle of a suboptimal BP and clearly correlates with crucial changes in neuronal functions linked to learning and memory.

Previously, we had demonstrated how strain-specific splicing of the *Camk2.1* gene in a marine insect controls the circadian timing of the species behaviour (Kaiser *et al.*, 2016). Mammalian *CAMK2B* is predominantly involved in the regulation of synaptic plasticity, and previous studies hinted at functional implications of alternative splicing of this gene (Bayer *et al.*, 2002; Bhattacharyya *et al.*, 2020; Brocke *et al.*, 1995; GuptaRoy *et al.*, 2000; O’Leary *et al.*, 2006; Sloutsky and Stratton, 2021). Our results show how the primate-specific weakening of a BP motif in the *CAMK2B* gene leads to primate-specific exon skipping and the generation of several primate-specific protein isoforms. In line with previous studies (Barbosa-Morais *et al.*, 2012; Gao *et al.*, 2015), changes in a *cis*-acting element, rather than the *trans*-acting environment, control the observed species-specific splicing differences. Interestingly, rather than affecting auxiliary enhancer or repressor sequences, the identified genomic differences specifically modulate the BP sequence, one of the canonical splicing motifs. Thus, alteration of BP strength can contribute to the decoupling of alternative splicing from changes in the *trans*-acting environment. As the *trans*-acting environment differs between different organs and tissues, our findings provide an explanation for species-specific splicing patterns that are present throughout different organs.

The BP is a prime target for what has been termed “evolutionary tinkering” (Jacob, 1977; Ule and Blencowe, 2019), meaning the gradual accumulation of mutations to promote new functions, while minimizing disruptive effects on existing functions. Introns often contain multiple functional BPs, leading to flexibility regarding BP selection, which may facilitate evolutionary adaptation of individual BPs; in addition, introns can be removed in multiple steps, a process termed recursive splicing (Wan et al., 2021). Our data suggest that evolutionary weakening of core splicing elements, including the BP, is a general principle to globally control species-specific alternative splicing. When comparing orthologous exons in humans and mouse, both species feature weaker BP motifs and splice sites in exons that are exclusively alternative in the respective species.

We also show that primate-specific CaMKIIβ protein isoforms subtly differ in their kinetic properties and in their substrate spectra. Kinetic differences are presumably mediated by conformational differences of various inactive and active states of the holoenzyme. However, the exact nature of these states is still under debate (Buonarati et al., 2021; Chao *et al.*, 2011; Myers et al., 2017; Sloutsky *et al.*, 2020). In our study, we confirm recent results that under steady-state conditions, the variable linker segment that is modulated by alternative splicing does not affect the cooperativity of the enzyme (Sloutsky *et al.*, 2020), but modulates the maximal activity at optimal calmodulin concentrations. While the observed effects are comparatively small, CaMKII isoforms represent the most abundant proteins at the post-synapse (Cheng *et al.*, 2006; Erondu and Kennedy, 1985), such that even small kinetic differences may translate into a large overall effect *in vivo*.

Similar to a previous publication (Bhattacharyya *et al.*, 2020), we find isoform-specific differences in CaMKII autophosphorylation in our analog-sensitive kinase assay. Exon 13-exclusion isoforms show a downregulation of the inhibitory autophosphorylations (T306/307), which prevent re-association of calmodulin and thereby the full activation of the enzyme. The complementary activating autophosphorylation (T287) occurs for all isoforms, but to a smaller extend in exon 16-exclusion isoforms. These findings corroborate and extend a study of fly CaMKII that had revealed isoform-specific differences in substrate specificity with isolated proteins *in vitro* (GuptaRoy *et al.*, 2000), suggesting direct interactions of the linker segment with selected target proteins.

Additionally, we have identified many novel CaMKIIβ substrates (Supplementary data 2), often featuring tyrosine-phosphorylations, which have previously only been reported for an artificial CaMKII construct (Sugiyama et al., 2008). While isoform- and group-exclusive phosphorylation targets exist, the isoforms also target many overlapping sites. An additional difference between the substrate spectra lies in the relative abundance of the various phosphorylation sites, showing that a given substrate is phosphorylated by different CaMKIIβ isoforms with a different probability. Being identical in the CaMKIIβ variants, the kinase domains *per se* cannot be the source of these differences. However, the flexible linker, modulated by alternative splicing, can change the probability that a particular substrate comes in contact with the active center.

CaMKIIβ also plays a structural role in synapses. Alternative splicing changes the affinity of the resulting isoforms to actin (O’Leary *et al.*, 2006) and thus the architecture of the cytoskeleton (Hoffman et al., 2013) and presumably of other protein networks, such as the PSD. It is therefore likely that the length and composition of the variable linker affect the positioning of CaMKIIβ isoforms within these structures and hence the exposure to specific substrates. Although CaMKIIβ readily dissociates from actin filaments after stimulation (Lin and Redmond, 2008; Shen and Meyer, 1999) it has been proposed that due to the transient nature of neuronal signaling, every CaMKII subunit only phosphorylates a single substrate during an individual calcium spike (Bayer and Schulman, 2019), emphasizing the impact of initial differences in subcellular localization.

As is expected for an enzyme involved in regulating synaptic plasticity, we observe a strong impact on long-term potentiation (LTP) in our mouse model of humanized *CAMK2B* alternative splicing, where the normal balance of splice isoforms is disrupted. Mechanistically, the unique functionality of the primate-specific splice variants could be based on any of the observed differences in molecular characteristics (or combinations thereof), such as changes in enzymatic activity, changes in CaMKIIβ autophosphorylation patterns, up- or down-regulation of specific phosphorylation targets or changes in sub-synaptic localization. For instance, the activating autophosphorylation T287, which is downregulated in the primate-specific exon 16-exclusion variants, has been implicated in decoding the frequency of calcium spikes during neuronal activity (De Koninck and Schulman, 1998; Hanson et al., 1994; Meyer *et al.*, 1992). Both of our mouse model strains, although having a slightly different genotype, show an identical phenotype with respect to transcriptome changes as well as LTP. This observation underscores a direct causal link between differences in *CAMK2B* alternative splicing and functional consequences for synaptic plasticity. Interestingly, our mouse model shows that a straight forward correlation of higher *CAMK2B* alternative splicing complexity with stronger LTP does not exist. The situation appears to be more complex and likely involves additional species-specific adaptations that then, together with *CAMK2B* alternative splicing, impact on LTP and learning and memory. Deciphering the coevolution of such networks, including species-specific changes in alternative splicing, will be a major challenge for future work but promises to shed light on the generation of species-specific cognitive abilities (Konopka et al., 2012; Wunderlich et al., 2014).

Our work provides, to our knowledge, the first example of a mouse model in which only a *cis*-acting element has been mutated to generate species-specific differences in alternative splicing, an approach which holds great promise in deciphering the exact mechanistic framework of splicing regulation and functional consequences. The observed effect on exon 16 splicing can potentially also be induced by other mutations, including single-nucleotide polymorphisms (SNPs) in the splice sites. We and others have also shown that modulation of other alternative exons, such as exon 13, affects CaMKIIβ functionality. We thus expect that SNPs or other mutations that modulate *CAMK2B*, and potentially *CAMK2A*, alternative splicing, alter *CAMK2* functionality and impact on a variety of neurological diseases.

Taken together, we connect evolutionary weakening of core splicing elements with species-specific alternative splicing and present a mouse model that connects primate specific *CAMK2B* alternative splicing with LTP, suggesting a prominent role of alternative splicing in the generation of species-specific cognitive abilities.

## Acknowledgment

The authors would like to thank Iva Lucic and Andrew Plested for discussions regarding CaMKII functionality, Kevan Shokat for insights into the analog-sensitive kinase system and members of the Heyd and Wahl labs for discussing all aspects of the project. Matthis Jahnel performed initial bioinformatic analysis to identify orthologous exons. AF was funded by a PhD Fellowship of the Boehringer Ingelheim Funds (BIF). Initial funding was provided by FU research funding. This study was supported by a grant from the Deutsche Forschungsgemeinschaft (TRR186/A15; project number 278001972) to FH and MCW.

## Author contribution

AF performed most experiments in this work with help from IW, ND and FS. AS, LMV and AV performed the electrophysiological characterization of the mouse model. RK generated the mouse model. AF, MP and AN analyzed the RNA-Seq data. YJ and BK measured the mass spectrometry samples and helped with data analysis. FH, MCW and AF designed the study, planned experiments, analyzed data, and wrote the manuscript with help from HU and DS. FH and MCW conceived and supervised the work.

## Material and Methods

### Materials availability statement

Material generated in this study is available upon reasonable request by email to FH.

#### Identification of endogenous *Camk2* alternative splicing isoforms

RT-PCR was performed with total RNA from human cerebellum (Clonetech, Cat# 636535), mouse cerebellum frog brain tissue and rhesus macaque total brain RNA (Zyagen, Cat# UR-201). Human cerebellum RNA contained material pooled from three male Asians, aged 21-29 (information provided by supplier). Total cerebellum RNA from mouse (*Mus musculus*) and total brain tissue RNA from frog (*Xenopus laevis*) was extracted via Trizol (see below). Human and macaque RNA was adjusted to 125 ng/μl, mouse and frog RNA to 500 ng/μl. Where necessary, specificity for *CAMK2B* was inferred by a gene-specific RT-primer, annealing to the less conserved exon 25 (numbering based on scheme in Figure 1A, human: TTG TGG TTG TCG TCG TCA TC; mouse: ACG AGG CAG ACA CAA ACA TG). Primers for the splice-sensitive radioactive PCR annealed to exons 9 and 23 (human/macaque for: CTC CAC GGT AGC ATC CAT GA; rev: AGT CCA TCC CTT CAA CCA GG; mouse for: CCA CCG TGG CCT CTA TGA T; rev: AAT CCA TCC CTT CGA CCA GG; Xenopus for: CCA CTG TTG CTT CCA TGA TG; rev: CCT GGT AGA AGG GAT AGA CT). PCR products were sequenced using the CloneJET PCR Cloning Kit (Thermo Fisher Scientific). RT-PCRs with radioactively labeled forward primers and quantification of PCR products were performed as previously described (Preussner et al., 2017).

#### Minigene Design and Splicing Assays

Minigenes were designed using the pcDNA3.1+ vector backbone. The minigenes contained the following sequences: Exon 11 with 300 bp of the downstream intron, Exon 16 with 100 bp of the upstream intron, the full intron in between exon 16 and 17, exon 17 with 300 bp of the downstream intron and exon 23 with 100 bp of the upstream exon (see Figure S1B).Alternative splicing of the minigenes was analyzed in N2A, SH-SY5Y, HEK and HeLa cells in biological triplicates. HEK and HeLa cells were cultivated in DMEM High Glucose medium (Biowest) with 10% fetal bovine serum (FBS) and penicillin/streptomycin (Biowest). N2A cells were cultivated in a 1:1 mix of Opti-MEM and DMEM medium (Opti-MEM with GlutaMAX, Gibco and DMEM with GlutaMAX, Gibco). SH-SY5Y cells were cultivated in DMEM High Glucose medium with 10% FBS, penicillin/ streptomycin and additional L-glutamine (1% v/v of 200 mM). Cells were seeded in 12-well plates with a concentration of 1*10^5^ cells/well (HEK, SH-SY5Y, N2A) or 1.5*10^5^ cells/well (HeLa). After 24h, the cells were transfected with 1 μg plasmid and 2 μl Roti-Fect (Carl Roth GmBH) transfection reagent per well. Cells were harvested 48 hours after transfection and RNA was extracted using RNA Tri-Liquid (BioSell) reagent according to the manufacture’s instruction. DNase I (Epicentre) digestion was performed according to the manufacture’s instruction to minimize contamination with plasmid DNA. Alternative splicing was analyzed by radioactive RT-PCR as described above, with a vector specific primer pair (T7f: TAATACGACTCACTATAGGG, BGHr: CCTCGACTGTGCCTTCTA).

#### siRNA Knockdown

Potential *trans*-acting factors were predicted with DeepClip (Grønning *et al.*, 2020). Additional *trans*-acting factors were predicted using RBP map (Paz *et al.*, 2014) and targeted predictions (Ray et al., 2013; Galarneau and Richard, 2005; García-Blanco et al., 1989; Rossbach et al., 2009). Knock downs were performed as described (Preussner et al, 2017) in HEK or N2A cells. Experiments were performed twice in biological triplicates.

#### Prediction of branch point sequences

The SVM-BPfinder (Corvelo *et al.*, 2010) tool was used to predict BP sequences. If not otherwise specified, human was selected as target organism to predict BP strength.

#### Expression and purification of selected CaMKIIβ isoforms

Selected CaMKIIβ isoforms were expressed in High Five insect cells via the baculovirus system. All purification steps were performed at 4°C. Cell pellets were resuspended in CaMKII lysis buffer (10 mM Tris/HCl pH7.5, 500 mM NaCl, 1 mM EDTA, 1 mM EGTA, 5% Glycerol, 1 mM DTT) supplemented with protease inhibitors (cOmplete, Roche) and lysed by sonication. Insoluble particles were separated by centrifugation at 21,500 rpm for 1 h. The soluble fraction was incubated with Strep-Tactin Sepharose beads (IBA Lifesciences) for 1 h and washed with CaMKII lysis buffer. Bound protein was eluted with CaMKII SEC buffer (50 mM PIPES pH 7.5, 500 mM NaCl, 1 mM EGTA, 10% glycerol, 1 mM DTT) containing 2.5 mM desthiobiotin (IBA Lifesciences). Eluted protein was concentrated and run on a Superose 6 10/300 GL size exclusion column (Cytiva) with CaMKII SEC buffer. Fractions were pooled according to SDS-PAGE and chromatogram, concentrated to approx. 1 mg/ml and flash frozen in single-use aliquots in liquid nitrogen. Before use, aliquots were thawed on ice, gently mixed by pipetting and centrifuged at 20,000 rcf for 5 min. Exactly equal concentrations were determined by repeated SDS-PAGE, Coomassie staining and quantification with ImageQuant TL (Cytiva).

#### Expression and purification of CaMKII substrate Syntide 2-GST

The sequence for Syntide 2 (PLARTLSVAGLPGKK) was expressed as a GST fusion protein, with a TEV cleavable N-terminal His-tag. A short linker (GGGGSGGGGS) was inserted between the Syntide 2 sequence and the C-terminal GST-tag. The fusion protein was expressed in BL21-RIL cells using auto-induction medium. All purification steps were performed at 4°C. Cell pellets were resuspended in lysis buffer (50 mM Tris-HCl pH 7.5, 150 mM NaCl, 20 mM imidazole, 1 mM DTT) containing protease inhibitors (cOmplete, Roche) and lysed by sonication. Insoluble particles were separated by centrifugation at 21,500 rpm for 1 h. The soluble fraction was loaded on a HisTrap Crude column (Cytiva) and eluted with a linear gradient of elution buffer (20 mM Tris-HCl pH 7.5, 300 mM NaCl, 500 mM imidazole, 1 mM DTT). Target fractions were pooled, supplied with TEV protease (self-made) and dialyzed against lysis buffer overnight. Digested samples were re-run on a HisTrap Crude column. The flow through was collected, concentrated and run on a High Load Superdex 75 26/60 size exclusion column (Cytiva), equilibrated with SEC buffer (20 mM PIPES pH 7.5, 50 mM NaCl). Target fractions were pooled, concentrated to 22 mg/ml and flash frozen in liquid nitrogen.

#### Expression and purification of human full-length tau (tau 441)

Human full-length tau (tau 441) was expressed as a fusion protein with an N-terminal His- and a C-terminal StrepII-tag. The protein was expressed in BL21 RIL cells in TB medium. Bacteria were grown at 37°C until an optical density of 0.6-0.8. Protein expression was induced with 1 mM IPTG for 3 h at 37°C. Cell pellets were resuspended in PBS buffer supplemented with 5 mM imidazole and protease inhibitors (cOmplete, Roche). Cells were lysed by sonication and incubated at 80°C in a water bath for 10 min with sporadic manual agitation. The lysate was cooled on ice for 10 min and supplemented with fresh protease inhibitors and 2 mM DTT. The lysate was cleared by centrifugation at 21,500 rpm for 30 min. The supernatant was loaded onto a HisTrap FF Crude 5 ml column (Cytiva) equilibrated with PBS supplemented with 5 mM imidazole and 1 mM DTT. The column was washed until baseline and the protein eluted with a linear gradient from 5-500 mM imidazole. Fractions were pooled based on the chromatogram and SDS-PAGE. Pooled fractions were loaded on a StrepTrap 5 ml column (Cytiva), equilibrated with PBS + 1 mM DTT. The column was washed until baseline and the protein was eluted with PBS containing 1 mM DTT and 2.5 mM desthiobiotin (IBA Lifesciences). Fractions were pooled based on the chromatogram and SDS-PAGE. The pooled sample was concentrated using a molecular weight cut-off of 3 kDa and run on a Superdex S200 26/60 (GE), equilibrated in PBS supplemented with 1 mM DTT. Fractions were pooled based on the chromatogram and SDS-PAGE, concentrated to ∼ 15 mg/ml and flash frozen in liquid nitrogen.

#### *In vitro* kinase assay

The protocol was adapted from (Coultrap and Bayer, 2012). CaMKII activity was measured by ^32^P incorporation into the substrate Syntide 2-GST or tau 441 (human). The model substrate Syntide 2 (Hashimoto and Soderling, 1987) was linked to GST to increase its molecular weight, facilitate purification and enable separation on an SDS-PAGE. Reactions were performed in 0.2 ml PCR-stripes. Purified CaMKIIβ was diluted to 10 nM in a mix containing 50 mM PIPES pH 7.2, 0.1% BSA, 2 mM CaCl_2_, 10 mM MgCl_2_, 50 μM Syntide 2-GST or 10 μM tau (human tau 441). The reaction was started by adding 1 nM to 4 μM calmodulin (Calbiochem) and 100 μM ATP (∼1 Ci mmol^-1^ [γ^32^P]-ATP). Reagents were pre-incubated at 30°C for 5 min. Reactions were carried out in a final volume of 30 μl for 2 min at 30°C. Reactions were terminated by adding 10 μl SDS sample buffer. Samples were run on a 12.5% SDS-PAGE, dried and analyzed via a photostimulable phosphor plate. Gels were quantified using ImageQuant 5.2 or ImageQuant TL (Cytiva). Results were plotted using Graph Pad Prism 6 and fit to a Hill equation (allosteric sigmoidal nonlinear fit). For the standard IVK assay, the experiment was repeated two times in triplicates. To compare the maximal activity at optimal calmodulin concentrations, V/Vmax_FL_ values for calmodulin concentrations from 100-1000 nM were pooled and plotted using Graph Pad Prism 6. Normal distribution and equality of variances was tested via Shapiro-Wilk test, Q-Q-Plots and F-test. Based on the results, a Student’s t-test or Welch’s t-test was performed. Resulting p-values were adjusted for multiple comparison using Holm’s method. Statistical analysis was performed in R and RStudio. For the autoactivity assay, the activation of CaMKII with varying concentrations of calmodulin was performed in the absence of the substrate protein. After a 2-minute incubation, the activation was quenched by addition of 5.3 mM EGTA. The substrate protein was added together with 3.3 mM MgCl_2_ to enable the phosphorylation reaction. The sample was incubated for 3 min and the reaction terminated with SDS sample buffer. The analysis was performed as described above.

#### Analog-sensitive kinase assay – pulldown and *in vitro* kinase assays

HEK cells were cultured as described above and seeded at a concentration of 0.2*10^5^ cells/ml and 12 ml in 10 cm dishes or 30 ml in 15 cm dishes. The CaMKIIβΔ13,16 analog-sensitive variant was transfected 24 h after seeding as described above, using 12 μg for 10 cm dishes and 36 μg for 15 cm dishes. 24 h after transfection, cells were harvested with trypsin, transferred to 1.5 ml reaction tubes and washed with PBS before being flash frozen in liquid nitrogen and stored at −80°C. Cell pellets corresponding to 3x 10 cm dishes and 2x 15 cm dishes were thawed on ice and resuspended in CaMKII lysis buffer (10 mM Tris/HCl pH7.5, 500 mM NaCl, 1 mM EDTA, 1 mM EGTA, 5% Glycerol, 1 mM DTT) supplemented with protease inhibitors (cOmplete EDTA-free, Roche). Cells were lysed by sonication on ice at 40% amplitude, 0.5 cycle and six rounds of 5 s. Lysates were cleared by centrifugation at 20,000 rcf, 4°C for 30 min. The supernatant was transferred to a new reaction tube and mixed with 50 μl pre-equilibrated StrepTactinXT beads (IBA) and supplemented with biotin-blocking solution (IBA). Samples were incubated for 1 h at 4°C with slow rotation. Beads were sedimented by centrifugation at 500 rcf, 4°C for 5 min. Beads were washed three times in CaMKII SEC buffer (50 mM PIPES pH 7.5, 500 mM NaCl, 1 mM EGTA, 10% glycerol, 1 mM DTT) and bound protein eluted CaMKII SEC buffer supplemented with 50 mM biotin (IBA). The eluate was dispersed into single-use aliquots, flash frozen in liquid nitrogen and stored at −80°C. To compare the AS variant to the wt kinase, a standard *in vitro* kinase assay (IVK) was performed as described above, using a limited range of calmodulin concentrations and roughly estimating the concentration via UV-absorption at 280 nm. To test the inhibition by various ATP analogs, the standard IVK assay was modified and set to a single calmodulin concentration of 100 nM. The reaction mixture contained varying concentrations of non-radioactive ATP (0-1 mM) or 0.5 mM of one of the following non-radioactive ATP analogs: N^6^-methyl-ATP, N^6^-etheno-ATP, N^6^-phenyl-ATP, N^6^-benzyl-ATP (Jena Bioscience).

#### Analog-sensitive kinase assay – *in vivo* labelling and thiophosphate enrichment

The thiophosphate enrichment strategy was based on (Michowski et al., 2020), with modifications. The analog-sensitive kinase variants were PCR amplified with primers omitting the Twin-Strep-tag and cloned back into the pcDNA3.1 expression plasmid, to avoid interference from the affinity tag. N2a cells were cultured as described above and seeded into 15 cm dishes at a concentration of 0.1*10^6^ cells/ml and 30 ml/dish. Cells were incubated for 24 h and transfected with 37.5 μg DNA and 75 μl Rotifect (Carl Roth) per 15 cm dish, as described above. Cells were grown for 48 h, washed with 20 ml PBS and subsequently 20 ml AS lysis buffer (20 mM PIPES pH 7.5, 150 mM NaCl, 10 mM MgCl_2_, 1 mM EGTA). The liquid was removed and the dish was carefully washed with 1.2 ml AS lysis buffer, supplemented with protease inhibitors (cOmplete, Roche), phosphatase inhibitors (PhosSTOP, Roche) and 0.5 mM TCEP. The liquid was removed thoroughly, the cells detached with a cell scraper, transferred to a reaction tube and kept on ice until all samples had been harvested. From then on, samples were processed in parallel in Protein LoBind tubes (ThermoFisher). Each 15 cm dish resulted in approximately 1.2 ml cell suspension, which was split into two 600 μl aliquots. The remaining cells were discarded. Each aliquot was supplemented with 75 μl detergent mix (3.6% nOG, 36 mM CaCl_2_) and briefly mixed. The labelling reaction was started by addition of 225 μl reaction mix (200 nM calmodulin, 0.1 mM N^6^-benzyl-ATPγS, 0.2 mM ATP, 3 mM GTP, PhosSTOP phosphatase inhibitors in AS lysis buffer) and incubated for 30 min at 30°C with sporadic manual agitation. For the untransfected control, calmodulin was omitted. The reaction was terminated by addition of EDTA/EGTA to a final concentration of 10 mM each. Labelled samples were briefly sonicated to create a homogeneous suspension and concentrations were determined by Pierce 660 nM assay (ThermoFisher). Samples were flash frozen in liquid nitrogen and stored at −80°C. For the western blot, aliquots were alkylated with 50 mM PNBM (p-nitrobenzyl mesylate, Agilent) at a final concentration of 2.5 mM for 1 h at RT. The reaction was terminated by addition of SDS sample buffer (containing DTT) and the samples analyzed via standard SDS-PAGE and semi-dry western blotting. The blot was developed using an anti-thiophosphate ester antibody (ab92570, Abcam) and an HRP-linked anti-rabbit antibody (Cell Signaling Technologies).

For thiophosphate enrichment, samples were thawed and lysate corresponding to 6 mg protein was transferred into a 15 ml tube for protein precipitation. All samples were equalized in volume with AS lysis buffer and supplemented with 5 volumes of ice-cold methanol/chloroform mix (ration 4:1), followed by 3 volumes of ice-cold H_2_O. The samples were thoroughly mixed, incubated 10 min on ice and centrifuged for 20 min at 2000 rcf. The resulting pellet, located at the interface, was washed in 5 volumes ice-cold methanol and centrifuged for 20 min at 2000 rcf. The supernatant was removed and the pellet dried at RT. The dried pellet was resuspended in 800 μl denaturation buffer (100 mM NH_4_HCO_3_, 2 mM EDTA, 10 mM TCEP adjusted to pH 7-8, 8 M urea), adjusted to 6 M urea with H_2_O and incubated at 55°C for 1 h with agitation at 300 rpm. The sample was slowly cooled to RT for 10 min and diluted to 2 M urea with 50 mM NH_4_HCO_3_ in H_2_O. TCEP (pH adjusted to 7-8) was added to a final concentration of 10 mM. Trypsin (Trypsin, TPCK treated from bovine pancreas, Sigma) was added at a ratio of 1:20 (w/w, based on starting material) and the samples were digested overnight at 37°C. The following morning, 10 M NaOH was added to a final concentration of 0.08 mM and the digestion continued for 3 h. The digest was acidified with 2.5% trifluoroacetic acid (TFA) to a final concentration of 0.1% and a pH of ∼ 2.5. If required, more TFA was added to lower the pH. The digest was centrifuged for 3 min at 1400 rcf and the supernatant aliquoted to a new tube. SepPak Plus cartridges (Waters) were equilibrated by sequential washing with 10 ml 0.1% TFA/50% acetonitrile (in H_2_O) and 10 ml 0.1% TFA (in H_2_O). The sample was loaded by passing it through the cartridge 5 times. The cartridge was washed with 10 ml 0.1% TFA (in H_2_O). Bound peptides were eluted with 4 ml 80% acetonitrile/0.1% acetic acid and dried overnight in a vacuum centrifuge. SulfoLink beads (ThermoFisher) were transferred to a Protein LoBind tube and washed with 200 mM HEPES pH 7.0. Beads were incubated with 200 mM HEPES pH 7.0, 25 μg/ml BSA for 10 min at RT in the dark. Beads were sequentially washed with 200 mM HEPES pH 7.0 and 2 times with 4 M urea, 0.1 M Tris pH 8.8, 10 mM TCEP (pH of stock solution ∼ 2.5, this lowers the total pH to ∼ 8.0). The dried peptides were resuspended in 4 M urea, 0.1 M Tris pH 8.8, 10 mM TCEP and acidified to pH 5 with 5% (v/v) formic acid. The peptide solution was added to the equilibrated beads and rotated overnight at RT in the dark. The next day, the beads were centrifuged at 2000 rcf for 3 min and the supernatant discarded. The beads were washed sequentially with 4 M urea in 20 mM HEPES pH 7.0, H_2_O, 5 M NaCl, 50% acetonitrile in H_2_O and 5% (v/v) formic acid. Unused binding sites were blocked by incubation with a fresh solution of 10 mM DTT for 10 min in the dark. Bound peptides were eluted in three steps with a solution of 2 mg/ml Oxone (Potassium peroxymonosulfate, Sigma) in H_2_O. Eluates were pooled and desalted using SepPak Plus cartridges as described above. Samples were dried in a vacuum centrifuge and stored at −80°C.

To remove remaining contaminants, peptides were further purified with Styrene Divinyl Benzene (SDB) StageTips. StageTips were prepared by inserting the material into standard 200 μl pipet tips and washing sequentially with methanol, 80% acetonitrile in 0.1% formic acid and two steps of 0.1% formic acid in H_2_O. The resuspended samples were acidified with 10% formic acid to a final concentration of 1%. The samples were loaded and passed through the StageTips, followed by sequential washing with 0.1% formic acid in H_2_O and two rounds of 80% acetonitrile in 0.1% formic acid. Bound peptides were eluted with 5% NH_4_OH in 60% acetonitrile, split into two equal aliquots and dried in a vacuum centrifuge. Peptides were measured on an Orbitrap Q Exactive HF (Thermo Scientific) or an Orbitrap Exploris 480 (Thermo Scientific). MS raw data were analyzed using MaxQuant (Version 1.6.5.0) against the UniProt mouse reference proteome (downloaded in November 2021, mouse, 25,367 entries). Subsequent analysis was done in python (version 3.8.5, Anaconda distribution) using the packages pandas, numpy, matplotlib, seaborn, upsetplot, scipty, sklearn. Contaminants and reverse peptide hits were removed and the analysis restricted to phosphorylated peptides with a localization probability ≥ 0.75. The overlap between the two datasets was calculated using the unique phosphosite (protein/gene name + identity of phosphorylated residue) as an index. The intensity values of both datasets were normalized before merging, using the min-max normalization: 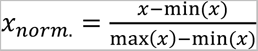.

Min(x) and max(x) were set to the respective minimal or maximal value of the individual datasets. When pooling replicates, an average intensity value was calculated. If only one replicate featured an intensity value for the respective target, this value was kept. Correlation matrices were calculated using a Pearson correlation coefficient.

The MS data have been deposited to the ProteomeXchange (Perez-Riverol et al., 2021) Consortium via the PRIDE partner repository with the dataset identifier PXD035346.

#### Generation of the mouse models

Mouse models were generated in the Transgenics Facility at the Max Delbrück Center for Molecular Medicine Berlin (MDC) under the supervision of Dr. Ralf Kühn. The models were based on the *CAMK2B* minigenes. CRISPR/Cas9 was used in C57Bl/6 mouse zygotes as described (Wefers et al., 2017) to remove the 100 bp initially found to harbor the *cis*-acting element in the endogenous mouse *Camk2β* gene. A synthetic gene was used as a repair template to insert the human ortholog of the excised sequence into the endogenous mouse gene (humanized strain). A deletion strain was generated in which the repair process failed and only the mouse sequence was deleted. Mice were handled according to institutional guidelines under experimentation licenses G0111/17-E65, T0100/03 and T0126/18 approved by the Landesamt für Gesundheit and Soziales (Berlin, Germany) and housed in standard cages in a specific pathogen-free facility on a 12 h light/dark cycle with ad libitum access to food and water.

#### RNA Seq analysis

##### Mouse model

Total RNA was extracted from mouse cerebellum tissue as described above (minigene splicing assay). 4 male wt, 2 male and 2 female heterozygous and 4 male homozygous mice of the humanized strain were selected for RNA-Seq. For library preparation, DNAse I-digested RNA samples were filtered using the polyA+ selection method at BGI Genomics and sequenced using DNBSeq PE150 sequencing. This yielded ∼50-60 million paired-end 150 nt reads. Reads were aligned to the GRCm38 genome using the STAR aligner (v.2.7.9a) (Dobin et al., 2013), yielding on average ∼75% uniquely mapped reads. Files were indexed using SAMtools (Danecek et al., 2021) and the splicing pattern analyzed using rMATS (v3.1.0) (Shen *et al.*, 2014). Downstream analyses and data visualization were performed using standard python code (v3.8.5). Data was visualized and sashimi plots generated via IGV (Robinson et al., 2011). Gene expression patterns were analyzed using Salmon (v1.8.0) (Patro et al., 2017) and DESeq2 (Love et al., 2014). Volcano plots were generated using GraphPad Prism 5-6. RNA-Sequencing data generated in this study are available under GEO #GSE208181.

##### Various mammals

Publicly available RNA-Seq data was analyzed for various mammals. For human, chimpanzee (*Pan trogodytes*), bonobo (*Pan paniscus*) and rhesus macaque (*Macaca mulatta*) data from cerebellum white tissue and cerebellum grey tissue from multiple individuals was selected. For gibbon (*Hylobates lar*), data from different brain regions from a single individual were selected. For gorilla (*Gorilla gorilla*) and orangutan (*Pongo pygmaeus*), data from cerebellum and total brain tissue was selected. For pig (*Sus scrofa*), data from cerebellum tissue was selected. Reads were aligned to the respective genome (human: GRCh38; chimpanzee: panTro6; bonobo: panPan1.1; gorilla: gorGor6; orangutan: ponAbe3 (*Pongo abelii*); Gibbon: nomLeu3 (*Nomascus leucogenys*); rhesus macaque: rheMac10; pig: SusScr11) using the STAR aligner (v.2.7.9a) (Dobin *et al.*, 2013). Subsequent analysis was performed as described above. To calculate %skipped values for *CAMK2B* exon 16, the sum of all individual exon 16 skipping events was calculated. For final visualization, cerebellum grey and white matter files (where available) were merged to create combined cerebellum files. A list of all used publicly available RNA-Seq data, including species, tissue, read length and used reference genome can be found in supplementary table S2.

#### Identification and analysis of orthogonal exons

Orthogonal exons in human and mouse were identified using the liftOver tool from the UCSC genome browser (Kent et al., 2002), with custom-optimization using the human genome assembly hg38 and the mouse genome assembly mm10. Publicly available RNA-Seq data from human brain tissue (47 samples from 35 individuals) and mouse brain tissue (9 samples from 9 individuals) were analyzed as described above and restricted to cassette exon events. Only splicing events supported by at least three datasets were kept. The results were filtered for a standard deviation of PSI (percent spliced in) below 0.2 and a minimal mean junction read count of 10. Alternative exons were defined as exons showing a PSI of < 0.9, and constitutive exons as exons showing a PSI of > 0.9. If orthologous alternative exons were identified in multiple transcripts, with different upstream or downstream exons, only the first listed entry was kept. Species-exclusive exons were defined as those being alternative in one, and constitutively included in the other species. If indicated, a further threshold of a minimal difference in PSI levels of 0.2 was applied. BP scores were calculated using SVMBP (Corvelo *et al.*, 2010) and splice site scores using MaxEntScan (Yeo and Burge, 2004). The difference between means was calculated using the paired Wilcoxon signed-rank test. To calculate a combined score of the BP motif and both splice sites, all parameter scores were normalized using the min-max normalization: 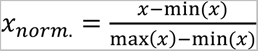 and a mean was calculated. Correlation between the PSI and various parameters was calculated using the Spearman singed-rank correlation coefficient.

#### Electrophysical characterization

All experiments regarding the electrophysical characterization were entirely performed in the research group of Prof. Dietmar Schmitz (Charité, NeuroCure) under the supervision of Dr. Alexander Stumpf. Hippocampal slices were prepared from adult C57/BL6J and transgenic (deletion and humanized) mice. Animals were anesthetized with isoflurane and decapitated. The brain was quickly removed and chilled in ice-cold sucrose-based artificial cerebrospinal fluid (sACSF) containing (in mM): NaCl 87, NaHCO_3_ 26, glucose 10, sucrose 50, KCl 2.5, NaH_2_PO4 1.25, CaCl_2_ 0.5 and MgCl_2_ 3, saturated with 95% (vol/vol) O_2_/5% (vol/vol) CO_2_, pH 7.4. Horizontal slices (300 μm) were cut and stored submerged in sACSF for 30 min at 35 °C and subsequently stored in ACSF containing (in mM): NaCl 119, NaHCO_3_ 26, glucose 10, KCl 2.5, NaH_2_PO4 1, CaCl_2_ 2.5 and MgCl_2_ 1.3 saturated with 95% (vol/vol) O_2_/5% (vol/vol) CO_2_, pH 7.4, at RT. Experiments were started 1 to 6 h after the preparation.

Recordings were performed in a submerged recording chamber (Warner instruments RC-27L), filled with ACSF with solution exchange speed set to 3-5 ml/min at RT (22-24°C). Stimulation electrodes were placed in the stratium radiatum of CA1 (near CA3) to stimulate Schaffer collaterals. Recording electrodes were placed in the str. radiatum of the CA1 field. Stimulation was applied every 10 s. In order to analyze the input-output relationship, stimulation intensities were adjusted to different FV amplitudes (0.05 mV increments, 0.05mV – 0.4 mV) and correlated with the corresponding field excitatory postsynaptic potential (fEPSP). Paired pulse ratios (PPR) were determined by dividing the amplitude of the second fEPSP (50 ms inter-stimulus interval) with the amplitude of the first (average of ten repetitions). Long term potentiation (LTP): Basal stimulation was applied every 10 s in order to monitor stability of the responses at least for 10 min before LTP was induced by one single high frequency stimulation train (100 pulses, 100 Hz). Magnitude of LTP was determined by normalizing the average of the initial fEPSP slopes 25-30 min and 55-60 min after LTP induction to average baseline fEPSP slope. Data collection and quantification was performed blindly. 1-way ANOVA and Dunnet’s-multi-comparison test was used to compare the mean LTP and PPR values of the transgenic animals (humanized and deletion) to the WT-control. LTP-induction by multiple high frequency trains was only performed in the humanized strain and in WT animals, thus an unpaired t-test was performed to compare these groups. Normal distribution of the data was tested via D-Agostino & Pearson omnibus normality test.

---

**Figure S1:**
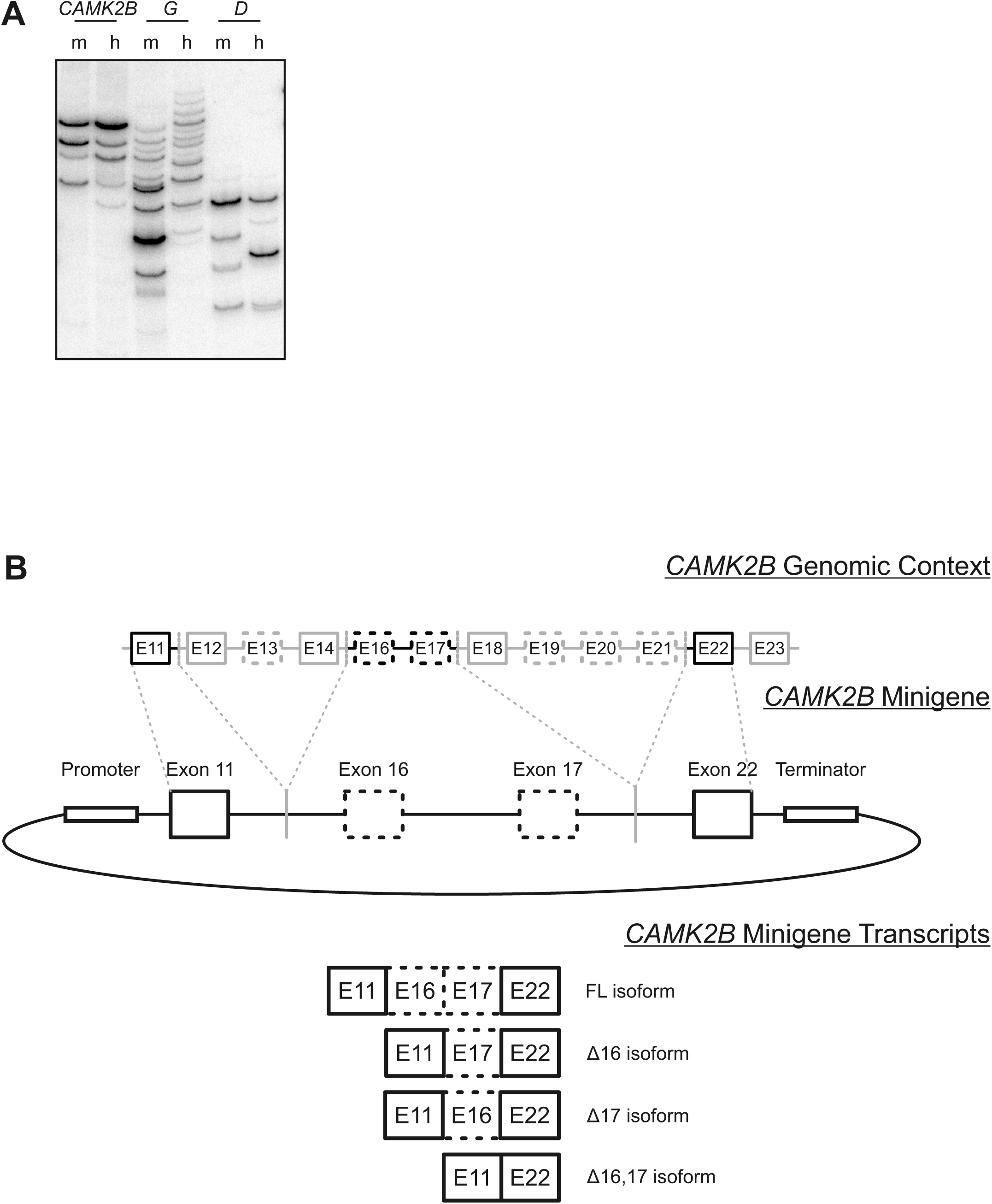
Endogenous *CAMK2B* splicing and minigene design. (A) Endogenous *CAMK2* splice isoforms were identified by radioactive isoform-specific RT-PCR using mouse (*Mus musculus*) and human cerebellum RNA. Isoforms were separated on a denaturing polyacrylamide gel and detected via autoradiography. m: mouse, h: human. (B) Schematic representation of the design for the *CAMK2B* minigenes. The top represents the genomic background, boxes represent exons, lines represent introns. Dashed boxes represent alternatively spliced exons. The middle part represents the minigene construct, containing two alternative exons flanked by two constitutive exons. Efficient transcription is ensured by promoter and terminator sequences. Dashed lines indicate which exonic and intronic regions from the genomic background were used in the minigene construct. The bottom displays all four possible alternatively spliced transcripts.

**Figure S2:**
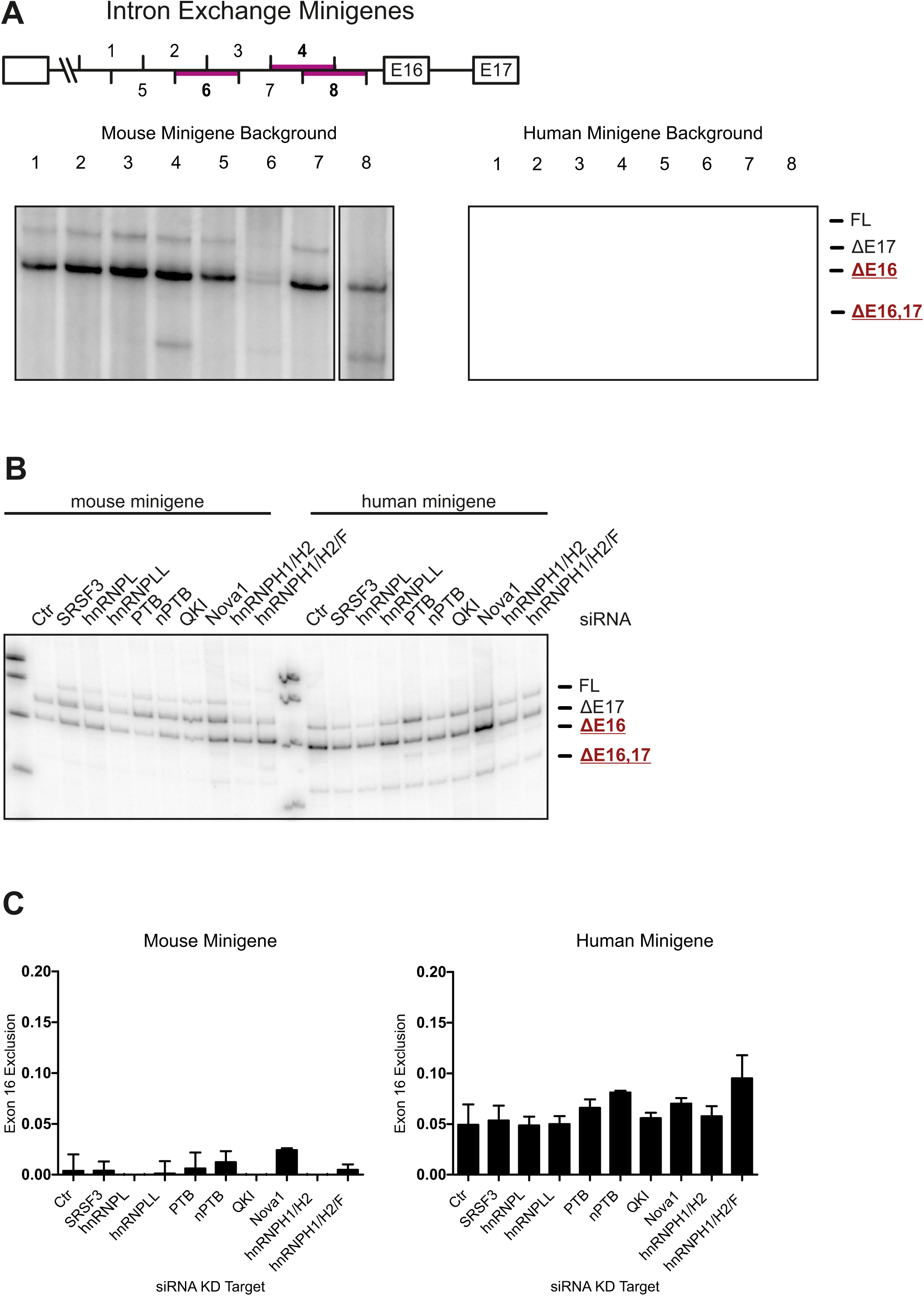
Identification of potential *trans*-acting factors. (A) Top: Schematic representation of the intron containing the identified functionally relevant *cis*-acting element. Numbers indicate 20 bp segments that were exchanged between the human and mouse construct. Purple lines highlight segments of functional relevance. Bottom: Human and mouse exchange minigenes were transfected into HEK cells and resulting splice isoforms identified by radioactive RT-PCR. (B) siRNA KD of potential *trans*-acting factors regulating the species-specific alternative splicing of *CAMK2B*. Indicated *trans*-acting factors were downregulated by siRNA-mediated KD in N2a cells. Human and mouse minigenes of *CAMK2B* were co-transfected and resulting splice isoforms identified by radioactive RT-PCR. Isoforms are indicated on the right and named according to the exons skipped. (C) Quantification of B, error bars indicate standard deviation (n=3).

**Figure S3:**
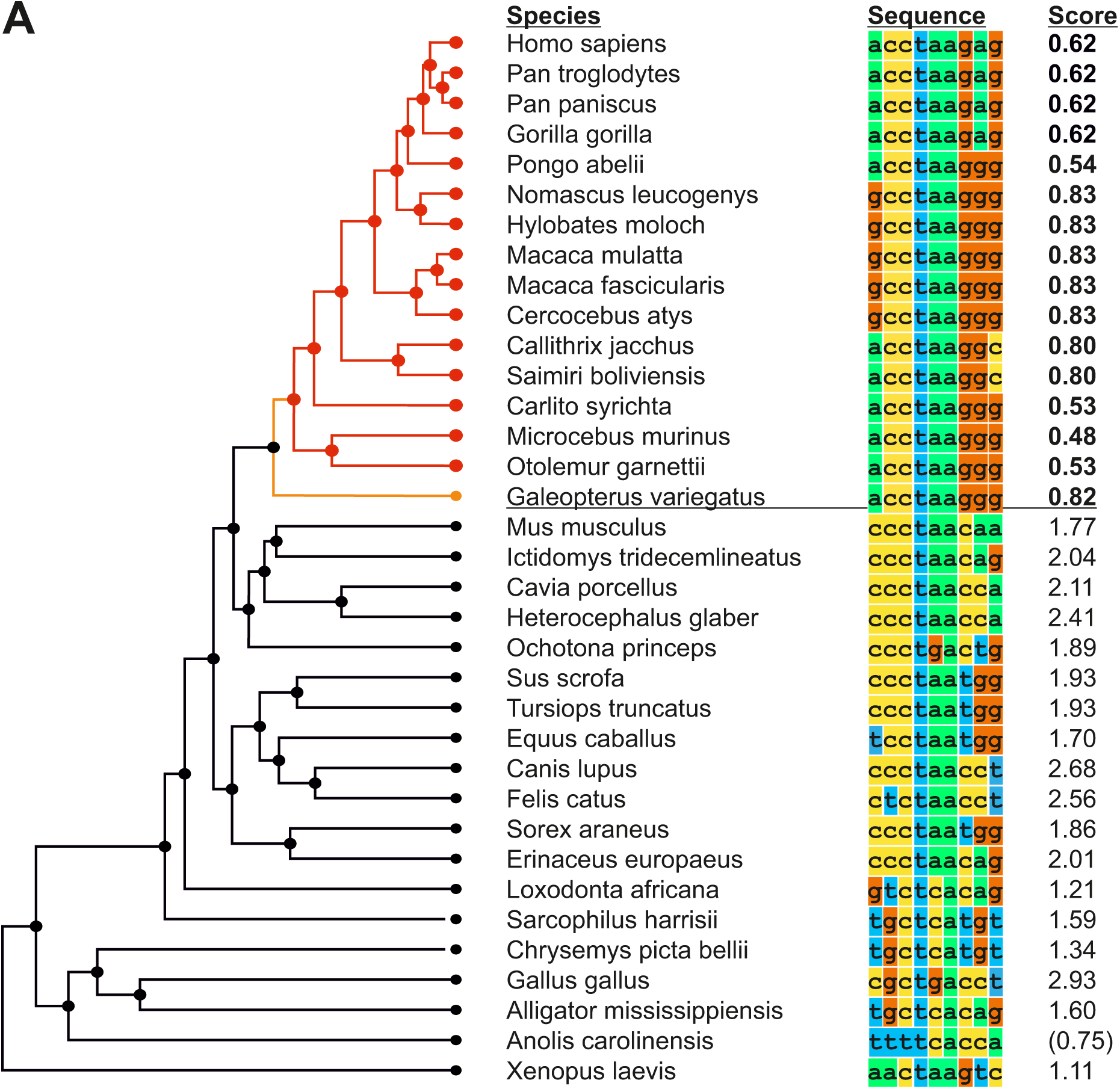
Evolution of the primate-specific branch point motif. (A) Alignment of the highest scoring BP sequence in various species. Only BPs within 100 nt of the 3’ splice site were considered. Left is a phylogenetic tree depicting evolutionary relationships. Red: Primate order, orange: Dermoptera order. Score refers to the BP motif score (scaled vector model) calculated via SVM-BPfinder (Corvelo *et al.*, 2010). Brackets indicate species for which no acceptable BP was found within 100 nt of the 3’ splice site.

**Figure S4:**
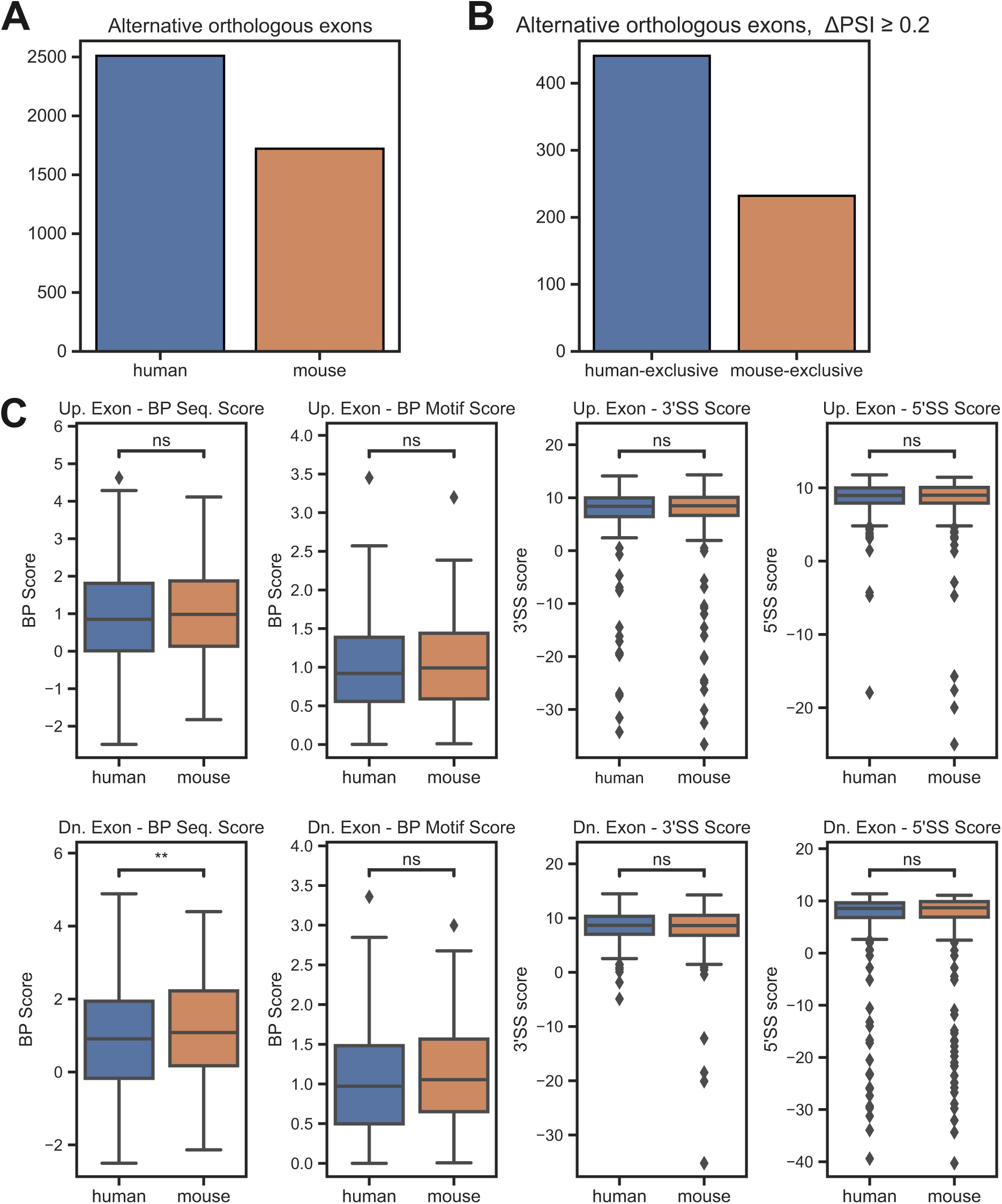
Branch point strength globally controls species-specific alternative splicing. (A) Number of alternative orthologous exons (PSI < 0.9) in human and mouse brain tissue. (B) Species-exclusive alternative orthologous exons. RNA-Seq data from different brain regions from mouse (n=47) and human (n=9) was analyzed to identify species-specific splicing pattern. The analysis was restricted to orthologous exons that are alternatively spliced in one species (PSI < 0.9) but not the other (PSI > 0.9) and show a minimal difference (ΔPSI) of 0.15. (C) Boxplot comparing human and mouse splicing element scores for constitutive exons upstream and downstream of human-exclusive alternative exons. PSI: percent spliced in, Up. Exon: exon upstream of the alternatively spliced exon of interest, Dn. Exon: exon downstream of the alternatively spliced exon of interest, BP Sequence Score: branch point sequence score, BP Motif Score: branch point motif score using a scaled vector model (Corvelo *et al.*, 2010), 3’/5’SS Score: splice site score (Yeo and Burge, 2004).

**Figure S5:**
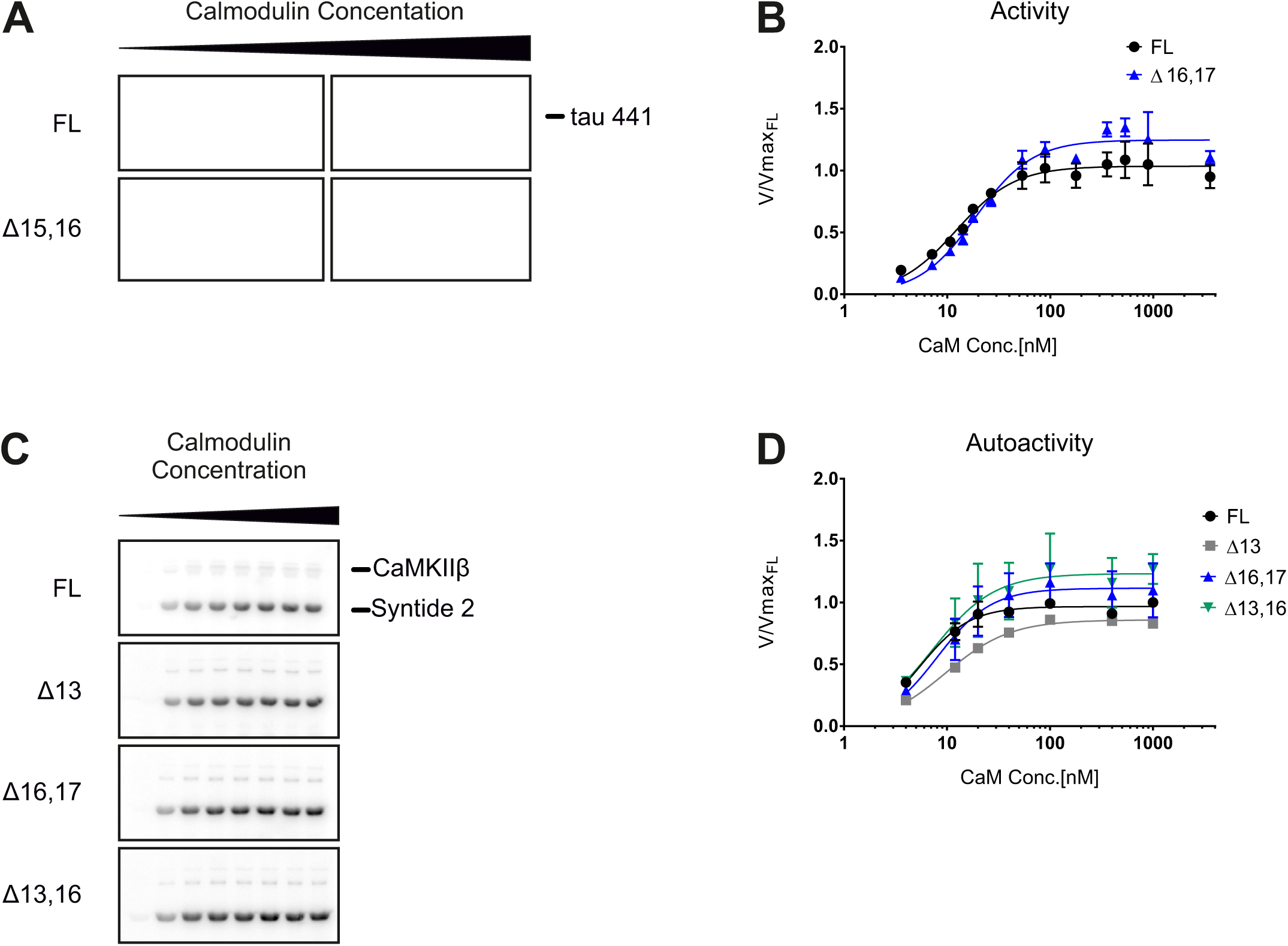
The kinetic differences are substrate independent. (A) *In vitro* kinase assay with different CaMKIIβ isoforms. CaMKII activity against a protein substrate (human full-length tau-441, with N-terminal polyhistidine- and C-terminal StrepII-tag) was measured as a function of calmodulin concentration. Direct phosphorylation of the substrate by CaMKIIβ was measured via ^32^P incorporation. Samples were separated on an SDS-PAGE and detected using autoradiography. (B) Quantification of A, normalized to the maximum activity of the FL isoform (n = 3). Data was fitted to a Hill equation (allosteric sigmoidal nonlinear fit) in GraphPad Prism 6. (C) *In vitro* kinase assay with different CaMKIIβ isoforms to test the autoactivity generated at increasing calmodulin concentrations. CaMKII was first activated with calmodulin in the presence of ATP. EGTA was added to chelate calcium and quench the binding of calmodulin. Addition of a protein substrate (Syntide 2, fused to GST) enabled detected of generated autoactivity via ^32^P incorporation. Samples were separated on an SDS-PAGE and detected using autoradiography. (D) Quantification of C, normalized to the maximum activity of the FL isoform (n=3)

**Figure S6:**
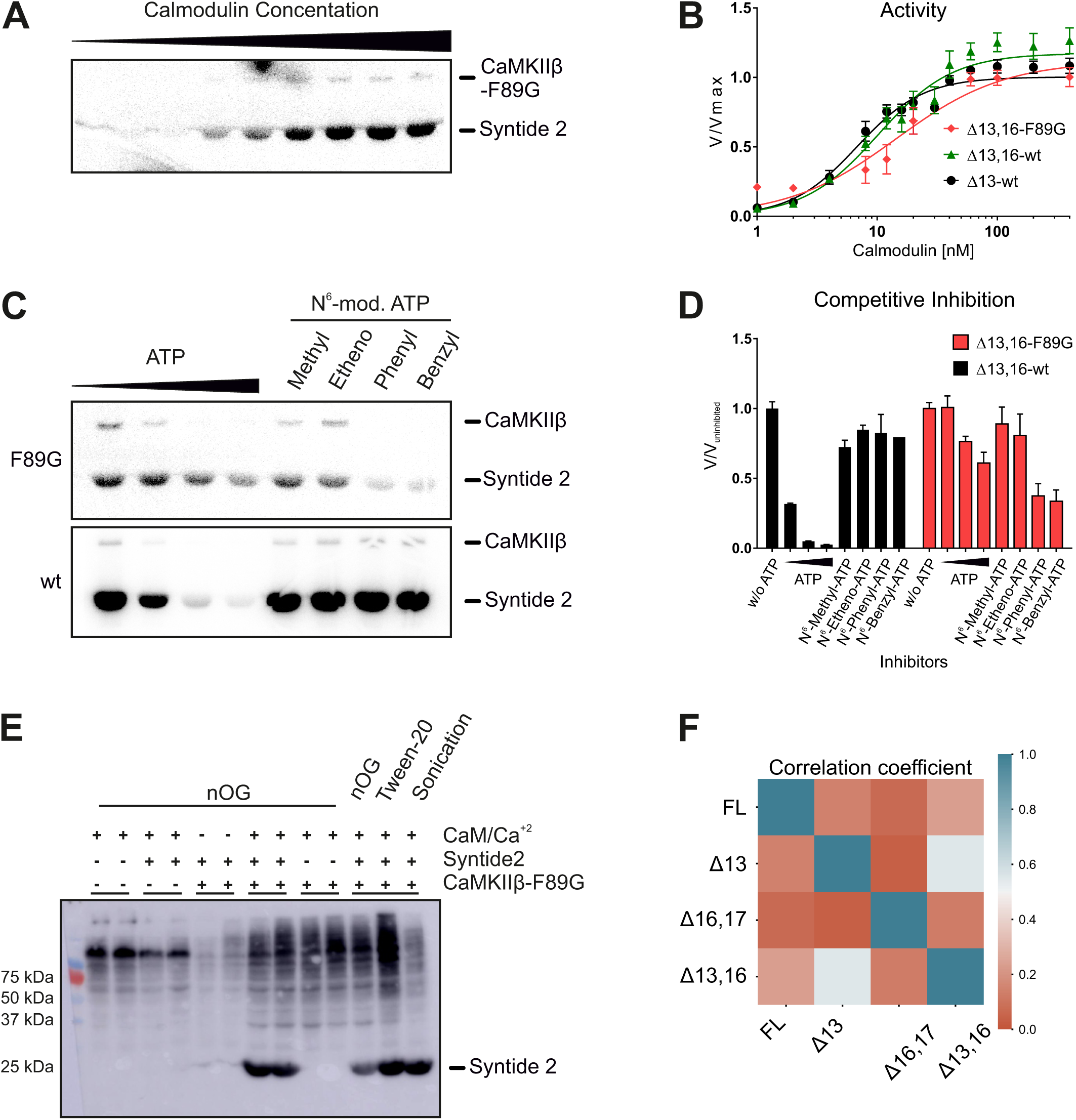
Analog-sensitive kinase assay. (A) *In vitro* kinase assay with the purified analog-sensitive (AS) variant of CaMKIIβ. CaMKII activity against a protein substrate (Syntide 2, fused to GST) was measured as a function of calmodulin concentration. Direct phosphorylation of the substrate by the analog-sensitive F89G variant was measured via ^32^P incorporation. Samples were separated on an SDS-PAGE and detected using autoradiography. (B) Quantification of A, normalized to the maximum activity at 400 nM calmodulin. Data for the wt variant of CaMKIIβΔ13 and CaMKIIβΔ13,16 are taken from Figure 5B, C. Error bars indicate standard deviation (n=3). (C) Inhibition of the AS variant of CaMKIIβ by various N^6^-modified ATP analogs. An *in vitro* kinase assay was performed at an optimal calmodulin concentration and supplemented with different non-radioactive ATP analogs. Enzymatic activity was measured as described for A. (D) Quantification of C, normalized to the non-inhibited signal (n=3). Note that the wt enzyme is not inhibited by N^6^-modified ATP analogs, whereas the AS variant is inhibited. (E) Western blot showing the labeling efficiency in permeabilized cells. N2a cells overexpressing Twin-Strep-CaMKIIIβΔ13,16-F89G were collected and permeabilized with nOG (n-octyl-β-D-glucopyranoside) or Tween-20 as indicated, or lysed by brief sonication. Reactions were performed with N^6^-benzyl-ATPγS in the presence/ absence of stimulating conditions (calmodulin/Ca^2+^) and/or an external substrate (Syntide 2, linked to GST). Samples were alkylated, run on an SDS-PAGE and analyzed via western blot with a thiophosphate ester-specific antibody. (F) Correlation matrix of the substrate spectra of different CaMKIIβ isoforms, as determined by an analog-sensitive kinase assay. The analysis was restricted to CaMKIIβ-specific targets. Additionally, CaMKIIβ autophosphorylation targets were removed from the analysis. A Person correlation coefficient was calculated based on the intensity values of individual phosphorylation sites.

**Figure S7:**
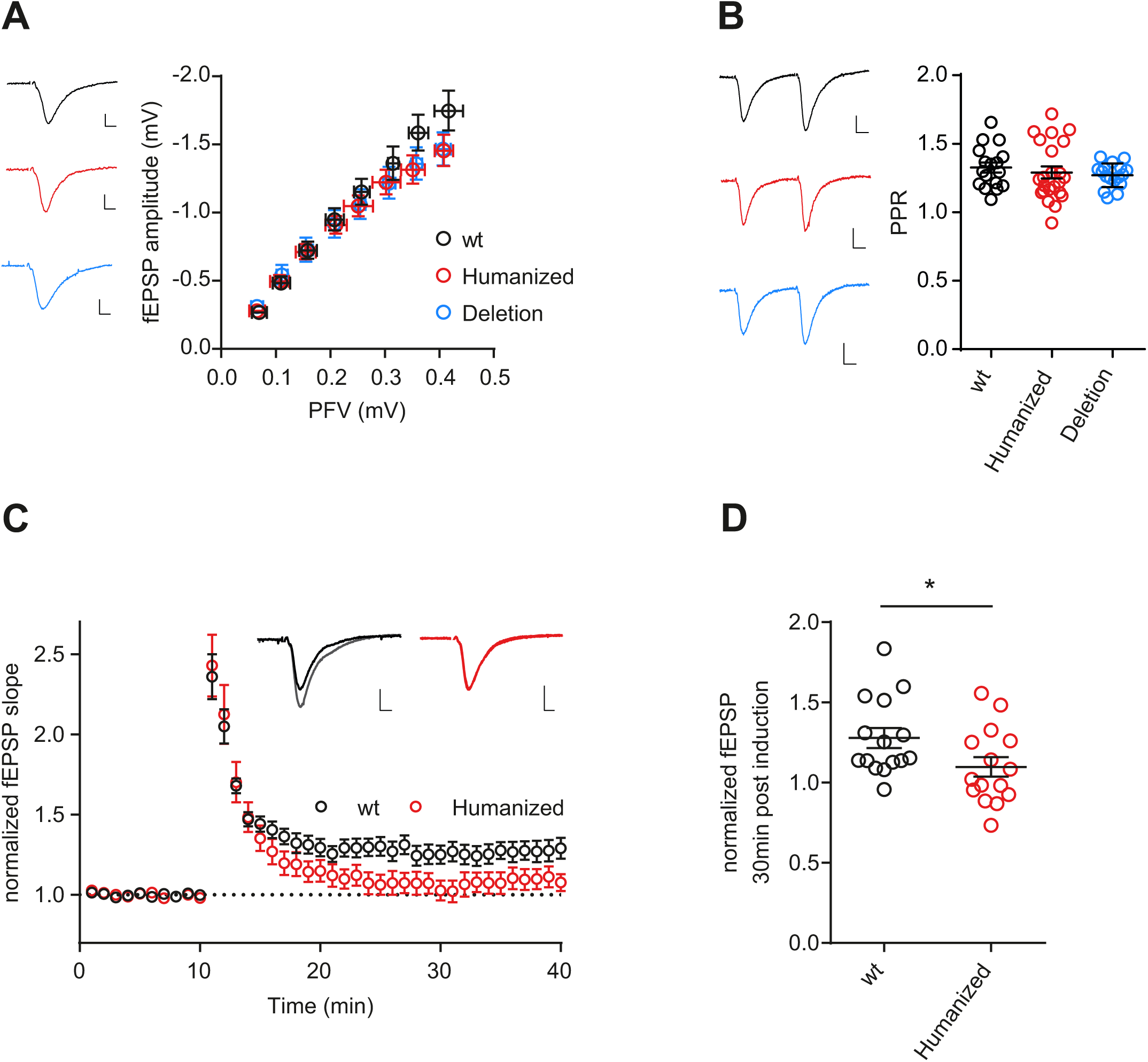
Electrophysiological characterization of the mouse model with humanized *Camk2β* splicing. (A) Input/ output characterization: relationship between amplitudes of presynaptic fiber volley (PFV) and field excitatory postsynaptic potential (fEPSP). Scale bars: 0.2 mV/ 5ms. (B) Short term plasticity (paired pulse ratio (PPR) with 50 ms inter-stimulus interval). Scale bars: 0.2 mV / 10ms. (C) Time course of LTP induction in CA3-CA1 synapses in acute hippocampal slices. LTP was induced after 10 min with four trains of 100 Hz, 1s. Example traces show average of baseline and potentiated field excitatory postsynaptic potentials (fEPSP) 30 min after LTP induction. Scale bar: 0.2 mV / 5 ms. wt (wild type): 15 slices, 6 mice, humanized (humanized strain, homozygote): 12 slices 6 mice. (D) Dot-plots depicting the field EPSP slope 30 min after LTP induction. *p<0.05, calculated by an unpaired t-test.

**Supplementary Table 1.** CaMKIIβ autophosphorylation sites are isoform-specific. The list of detected phosphorylation sites was restricted to gene names CaMK2a, CaMK2b, CaMK2d. Numbers are Log2 ratio between the average intensity values for this isoform vs. controls. Unique: this phosphorylation target was not detected in the corresponding control samples (UT and K43R) and thus no Log2 ratio could be calculated. Absent: this phosphorylation site was not detected in the corresponding sample.

| Unique phospho-site | Log2 (Enrichment) |  |  |  | Location of phosphosite | Function |
| --- | --- | --- | --- | --- | --- | --- |
|  | FL/Ctr | 13/Ctr | 16,17/Ctr | 13,16/Ctr |  |  |
| **Camk2b-S315** | unique | absent | unique | absent | Exon 12 | F-actin binding (Kim et al., 2015) |
| **Camk2b-T320** | unique | absent | unique | absent | Exon 13 | F-actin binding (Kim et al., 2015) |
| **Camk2b-T321** | unique | absent | unique | absent | Exon 13 | F-actin binding (Kim et al., 2015) |
| **Camk2a-T306** | unique | absent | unique | absent | Exon 12 | Inactivating (Colbran and Soderling, 1990) |
| **Camk2a-T307** | unique | unique | unique | unique | Exon 12 | Inactivating (Colbran and Soderling, 1990) |
| **Camk2b-S534** | 1,17 | 0,95 | -0,51 | -0,81 | Hub Domain | NA |
| **Camk2b-S280** | 5,68 | 5,58 | 4,58 | 4,98 | Exon 11 | GlcNAc site (Erickson et al., 2013) |
| **Camk2b-T287** | 6,57 | 6,48 | 5,47 | 5,88 | Exon 11 | Activating (Miller et al., 1988) |
| **Camk2d-S280** | 2,74 | 2,47 | 1,58 | absent | Exon 11 | GlcNAc site (Erickson et al., 2013) |
| **Camk2d-T287** | 2,74 | 2,47 | 1,58 | absent | Exon 11 | Activating |
| **Camk2b-T254** | unique | unique | unique | unique | Kinase Domain | NA |
| **Camk2b-S276** | unique | unique | absent | unique | Exon 11 | NA |
| **Camk2b-S71** | absent | unique | absent | unique | Kinase Domain | NA |

**Supplementary Table S2.** Publicly available RNA-Seq datasets used in this study.

| SRA | Species | Tissue | Read length | paired/single | Reference Genome |
| --- | --- | --- | --- | --- | --- |
| SRR8750487 | *Human* | Cerebellar White Matter | 150 bp | paired-end | GRCh38 |
| SRR8750488 | *Human* | Cerebellar White Matter | 150 bp | paired-end | GRCh38 |
| SRR8750489 | *Human* | Cerebellar White Matter | 150 bp | paired-end | GRCh38 |
| SRR8750490 | *Human* | Cerebellar White Matter | 150 bp | paired-end | GRCh38 |
| SRR8750491 | *Human* | Cerebellar Grey Matter | 150 bp | paired-end | GRCh38 |
| SRR8750492 | *Human* | Cerebellar Grey Matter | 150 bp | paired-end | GRCh38 |
| SRR8750493 | *Human* | Cerebellar Grey Matter | 150 bp | paired-end | GRCh38 |
| SRR8750647 | *Pan troglodytes* | Cerebellar White Matter | 150 bp | paired-end | panTro6 |
| SRR8750679 | *Pan troglodytes* | Cerebellar White Matter | 150 bp | paired-end | panTro6 |
| SRR8750711 | *Pan troglodytes* | Cerebellar White Matter | 150 bp | paired-end | panTro6 |
| SRR8750648 | *Pan troglodytes* | Cerebellar Grey Matter | 150 bp | paired-end | panTro6 |
| SRR8750680 | *Pan troglodytes* | Cerebellar Grey Matter | 150 bp | paired-end | panTro6 |
| SRR8750712 | *Pan troglodytes* | Cerebellar Grey Matter | 150 bp | paired-end | panTro6 |
| SRR8750448 | *Pan paniscus* | Cerebellar White Matter | 150 bp | paired-end | panPan1.1 |
| SRR8750449 | *Pan paniscus* | Cerebellar White Matter | 150 bp | paired-end | panPan1.1 |
| SRR8750450 | *Pan paniscus* | Cerebellar White Matter | 150 bp | paired-end | panPan1.1 |
| SRR8750451 | *Pan paniscus* | Cerebellar Grey Matter | 150 bp | paired-end | panPan1.1 |
| SRR8750452 | *Pan paniscus* | Cerebellar Grey Matter | 150 bp | paired-end | panPan1.1 |
| SRR8750595 | *Pan paniscus* | Cerebellar Grey Matter | 150 bp | paired-end | panPan1.1 |
| SRR5804509 | *Gorilla gorilla* | Cerebellum | 101 bp | paired-end | gorGor6 |
| SRR5804501 | *Gorilla gorilla* | Cerebellum | 101 bp | paired-end | gorGor6 |
| SRR306801 | *Gorilla gorilla* | Brain | 101 bp | paired-end | gorGor6 |
| SRR10393301 | *Pongo pygmaeus abelii* | Testis | 150 bp | paired-end | ponAbe3 |
| SRR10393303 | *Pongo pygmaeus abelii* | Testis | 150 bp | paired-end | ponAbe3 |
| SRR10393302 | *Pongo pygmaeus abelii* | Testis | 150 bp | paired-end | ponAbe3 |
| SRR10393304 | *Pongo pygmaeus abelii* | Testis | 150 bp | paired-end | ponAbe3 |
| DRR128395 | *Pongo pygmaeus* | Skin | 125 bp | paired-end | ponAbe3 |
| DRR128394 | *Pongo pygmaeus* | Skin | 100 bp | paired-end | ponAbe3 |
| DRR128393 | *Pongo pygmaeus* | Skin | 100 bp | paired-end | ponAbe3 |
| SRR306792 | *Pongo pygmaeus* | Brain | 150 bp | paired-end | ponAbe3 |
| SRR5804517 | *Hylobates lar* | Cerebellum | 100 bp | paired-end | nomLeu3<br>(*Nomascus leucogenys*) |
| SRR5804510 | *Hylobates lar* | Dorsolateral Prefrontal Cortex | 100 bp | paired-end | nomLeu3<br>(*Nomascus leucogenys*) |
| SRR5804511 | *Hylobates lar* | Ventrolateral Prefrontal Cortex | 100 bp | paired-end | nomLeu3<br>(*Nomascus leucogenys*) |
| SRR5804512 | *Hylobates lar* | Premotor Cortex | 100 bp | paired-end | nomLeu3<br>(*Nomascus leucogenys*) |
| SRR5804513 | *Hylobates lar* | Primary Visual Cortex | 100 bp | paired-end | nomLeu3<br>(*Nomascus leucogenys*) |
| SRR5804514 | *Hylobates lar* | Anterior Cingulate Cortex | 100 bp | paired-end | nomLeu3<br>(*Nomascus leucogenys*) |
| SRR5804515 | *Hylobates lar* | Striatum | 100 bp | paired-end | nomLeu3<br>(*Nomascus leucogenys*) |
| SRR5804516 | *Hylobates lar* | Hippocampus | 100 bp | paired-end | nomLeu3<br>(*Nomascus leucogenys*) |
| SRR8750552 | *Macaca mulatta* | Cerebellar White Matter | 150 bp | paired-end | rheMac10 |
| SRR8750553 | *Macaca mulatta* | Cerebellar White Matter | 150 bp | paired-end | rheMac10 |
| SRR8750554 | *Macaca mulatta* | Cerebellar White Matter | 150 bp | paired-end | rheMac10 |
| SRR8750549 | *Macaca mulatta* | Cerebellar Grey Matter | 150 bp | paired-end | rheMac10 |
| SRR8750550 | *Macaca mulatta* | Cerebellar Grey Matter | 150 bp | paired-end | rheMac10 |
| SRR8750551 | *Macaca mulatta* | Cerebellar Grey Matter | 150 bp | paired-end | rheMac10 |
| SRR11939284 | *Sus scrofa* | Cerebellum | 150 bp | paired-end | SusScr11 |
| SRR11939285 | *Sus scrofa* | Cerebellum | 150 bp | paired-end | SusScr11 |
| SRR11939286 | *Sus scrofa* | Cerebellum | 150 bp | paired-end | SusScr11 |

Supplementary data 1: Orthologs exons in mouse and human

Supplementary data 2: Mass spec data of analog sensitive kinase assay

